# Biases in ARG-based inference of historical population size in populations experiencing selection

**DOI:** 10.1101/2024.04.22.590609

**Authors:** Jacob I. Marsh, Parul Johri

**Affiliations:** Department of Biology, University of North Carolina, Chapel Hill, NC 27599; Department of Genetics, University of North Carolina, Chapel Hill, NC 27599; Integrative Program for Biological and Genome Sciences, University of North Carolina, Chapel Hill, NC 27599

**Keywords:** Demographic inference, ancestral recombination graph, background selection, selective sweeps, human population genomics, *Drosophila melanogaster*

## Abstract

Inferring the demographic history of populations provides fundamental insights into species dynamics and is essential for developing a null model to accurately study selective processes. However, background selection and selective sweeps can produce genomic signatures at linked sites that mimic or mask signals associated with historical population size change. While the theoretical biases introduced by the linked effects of selection have been well established, it is unclear whether ARG-based approaches to demographic inference in typical empirical analyses are susceptible to mis-inference due to these effects. To address this, we developed highly realistic forward simulations of human and *Drosophila melanogaster* populations, including empirically estimated variability of gene density, mutation rates, recombination rates, purifying and positive selection, across different historical demographic scenarios, to broadly assess the impact of selection on demographic inference using a genealogy-based approach. Our results indicate that the linked effects of selection minimally impact demographic inference for human populations, though it could cause mis-inference in populations with similar genome architecture and population parameters experiencing more frequent recurrent sweeps. We found that accurate demographic inference of *D. melanogaster* populations by ARG-based methods is compromised by the presence of pervasive background selection alone, leading to spurious inferences of recent population expansion which may be further worsened by recurrent sweeps, depending on the proportion and strength of beneficial mutations. Caution and additional testing with species-specific simulations are needed when inferring population history with non-human populations using ARG-based approaches to avoid mis-inference due to the linked effects of selection.

## Introduction

Inferring the demographic histories of natural populations using population-genetic approaches provides fundamental insights into both evolutionary and ecological species dynamics. Demographic inference methods predict the effective size of ancestral populations at different time points (epochs) in the past which best explains observed levels and patterns of neutral diversity among sequenced individuals (reviewed in Beichman et al. 2018; Marchi et al. 2021). Inferred population history can be correlated with archaeological and anthropological findings to, for example, uncover the date and scale of population bottlenecks and migration events (Mellars 2006; Jouganous et al. 2017; Bergström et al. 2020; Hu et al. 2023). Additionally, demographic history is needed to establish a null model for the inference of selection across the genome (Teshima et al. 2006; Caicedo et al. 2007; Nielsen et al. 2007; Thornton and Jensen 2007; Jensen et al. 2019). As we accumulate increasingly rich genomic resources and apply advanced methods for inferring evolutionary parameters of populations, it is essential that we understand the interactive effects between selection and historic demography on contemporary variation, including the ways in which selection can impact empirical inference of demography using modern methods.

Demographic history has traditionally been inferred using statistical summaries of genomic variation such as the site frequency spectrum (SFS; Gutenkunst et al. 2009; Kamm et al. 2020; Excoffier et al. 2021), linkage disequilibrium (LD; Hayes et al. 2003; Harris and Nielsen 2013; Ragsdale and Gutenkunst 2017), by approximate Bayesian computation approaches that use multiple summary statistics to perform inference (Beaumont et al. 2002; Duchen et al. 2013; Kapopoulou et al. 2018; Johri et al. 2020), or by modelling local genealogies (Li and Durbin 2011; Schiffels and Durbin 2014; Terhorst et al. 2017; Schiffels and Wang 2020). SFS-based approaches such as ∂*a*∂*i* (Gutenkunst et al. 2009) and *fastsimcoal2* (Excoffier et al. 2021) fit the observed SFS at putatively neutral sites to the expected SFS under a range of pre-determined demographic scenarios using diffusion theory and coalescent simulations respectively. However, SFS-based methods assume independence between segregating sites, which is violated when using dense variant data due to linkage disequilibrium between proximal loci. In contrast, sequentially Markovian coalescent (SMC) methods typically employ Hidden Markov Models (HMM) to generate local genealogies between historical recombination breakpoints across the genome, allowing for the estimation of genome-wide coalescence rates correlating to a specific branch-length across the genealogies (Li and Durbin 2011; Schiffels and Durbin 2014; Schiffels and Wang 2020). The coalescence rate across the genome is inversely proportional to the number of recombining individuals at a given time point in a panmictic population, thus the inverse instantaneous coalescent rate (IICR) can be used as a scaling factor to estimate relative effective population size even if it is also influenced by population structure and migration rate and should thus be interpreted with care (Mazet et al. 2016; Chikhi et al. 2018). SMC-based methods are limited to very few individuals (2-16 haplotypes) which limits analysis of recent demography. *SMC++* (Terhorst et al. 2017) was subsequently developed to address this limitation, accommodating the analysis of hundreds of samples to improve inference of recent demography. Furthermore, emerging methods for capturing detailed genealogies across larger populations present additional promising opportunities to improve demographic inference by harnessing richer sources of genetic information (Kelleher et al. 2019; Speidel et al. 2019; Zhang et al. 2023).

Ancestral recombination graphs (ARGs) capture detailed genealogical relationships between individuals by recording all ancestral recombination/coalescence events throughout the genome (Hudson 1991; Griffiths and Marjoram 1997), typically employing HMM algorithms similarly to SMC-based methods (Li and Stephens 2003). Among recently developed scalable ARG approximation tools, *Relate* has emerged as the most accessible and widely used for demographic inference as of yet (*e.g.* Speidel et al. 2019; Almarri et al. 2021; Minadakis et al. 2023; Moorjani and Hellenthal 2023; Teterina et al. 2023; Ishigohoka and Liedvogel 2024). To approximate an ARG, *Relate* estimates topological features of local genealogies (trees) based on position-specific distance matrices calculated between haplotypes. Following the estimation of local tree topology, *Relate* then estimates branch lengths using a Markov Chain Monte Carlo (MCMC) algorithm, initially assuming constant effective population size. Estimated branch lengths are then used to re-estimate historical population sizes from the inverse of the averaged coalescence rate at different epochs, allowing in turn for branch lengths to be re-estimated under the inferred demography. This process is iteratively repeated to co-estimate branch lengths and historical population size. The authors presented simulations of human data suggesting that *Relate* provides improved inference when compared to traditional local genealogy methods *MSMC* and *SMC++,* particularly for estimates of population size in the recent past (Speidel et al. 2019).

Although most demographic inference approaches perform accurately when tested with simulations assuming strict neutrality, indirect evolutionary forces, notably background selection (BGS) due to purifying selection (Charlesworth et al. 1993) and selective sweeps caused by positive selection (Maynard Smith and Haigh 1974), can produce genomic signatures that mimic or mask signals associated with historical population size change (Nicolaisen and Desai 2013), confounding inference of demography (Messer and Petrov 2013; Ewing and Jensen 2016; Schrider et al. 2016; Johri et al. 2021; Boitard et al. 2022; Smith and Hahn 2023). For example, a population bottleneck caused by non-selective factors (*e.g*., founder effects following migration) will result in a population having fewer immediate ancestors in subsequent generations; a hard selective sweep for a specific allele similarly leads to fewer immediate ancestors in subsequent generations due to increased fitness for a small proportion of the population (Teshima et al. 2006). Accurate inference of historical population sizes therefore becomes a pre-requisite for, and pre-requires, accurate inference of selection - presenting a circular dilemma (Johri, Aquadro, et al. 2022).

While a few recent simulation-based approaches have developed methods to jointly infer selection and demography while accounting for the linked effects of selection (Sheehan and Song 2016; Johri et al. 2020; Johri et al. 2023), the majority of widely-used demographic inference approaches assume strict neutrality, *i.e*., ignore the effects of direct and indirect selection, which is typically attempted by masking functional regions (*e.g*., conserved elements, protein-coding exons) from the analysis. However, LD with selected loci often extends beyond functional regions, violating assumptions of neutrality by impacting genetic diversity at putatively neutral sites. Findings by Pouyet *et al*. (2018) suggest that the majority of the human genome is influenced by BGS and biased gene conversion which in turn biases the SFS and demographic inference. Furthermore, Johri *et al*. (2021) demonstrated that BGS biases demographic inference when purifying selection is sufficiently pervasive, and spurious inference of recent population expansion is observed under population decline and equilibrium scenarios when using *fastsimcoal2* (Excoffier et al. 2021) and *MSMC* (Schiffels and Wang 2020). In contrast, Boitard *et al*. (2022), who explored the relative decrease in effective population size due to BGS and selective sweeps, suggest that the heterogeneity in the rate of coalescence across the genome, as generated by the linked effects of selection, can instead lead to spurious apparent decreases in effective population size. While previous studies clearly outline the theoretical biases introduced by BGS and selective sweeps to demographic inference, further analysis is needed to determine the extent to which the linked effects of selection impact empirical population genomic inference using ARG-based approaches where complex genome architecture and realistic effects of selection are present. Moreover, because we do not have precise estimates of the fraction and strength of new beneficial mutations in natural populations, it has been unclear if a reasonable extent of recurrent positive selection will significantly impact linked neutral variation in human and *D. melanogaster*-like populations.

Here, we test how demographic inference using an ARG-based approach is affected by selection in complex genomes using highly realistic simulations relevant to human and *Drosophila melanogaster*-like populations. To do so, we developed forward simulations incorporating variable gene density, recombination rates, and mutation rates, using realistic parameters for human and *D. melanogaster*-like chromosomes. We assess the effects on inference of different simulated demographic histories, with the effects of background selection and recurrent sweeps, parameterized by previously published estimates. Our findings provide insights into pitfalls that arise when ARG-based demographic inference approaches are applied to human populations and other species. Moreover, we collate previous estimates of positive selection parameters inferred from human and *D. melanogaster*-like populations, providing a potentially useful resource to other researchers (Table 1).

**Table 1.** Summary of empirically estimated positive selection parameters for different animal populations, including the selection inference method used. Confidence interval values provided where available in parentheses. Annotation type of selected sites used for analysis is provided: “CDS” is protein coding sites, “Nonsyn”, nonsynymous, “0-fold”, 0-fold degenerate coding sites.

| Taxon | Population | $f_{pos}$ | $2N_{anc}\bar{s}_a$ | $\alpha$ | Reference | Selected sites | Selection inference method |
| --- | --- | --- | --- | --- | --- | --- | --- |
| Human | African |  |  | 0.135 | Uricchio et al. 2018 | CDS | ABC-MK |
|  | YRI | 0.01 | 6.05 | 0.25 (0.24-0.252) | Zhen et al. 2021 | Nonsyn | Composite likelihood approach |
|  | YRI |  |  | 0.24 | Zhen et al. 2021 | Nonsyn | DFE-alpha (Eyre-Walker and Keightley 2009) |
|  | Mixed |  |  | 0.24 (0.18-0.29) | Galtier 2016 | Nonsyn | Grapes |
|  | Mixed |  |  | 0 (-0.30-0.24) | Eyre-Walker & Keightley 2009 | 0-fold | DFE-alpha |
|  | Mixed | 0.000023 (0.0000022-0.0076) | 0.0064 (0.0007-0.1084) |  | Castellano et al. 2019 | CDS | polyDFE v2.0 (Tataru and Bataillon 2019) using only the SFS |
| Chimpanzee | Western |  |  | -0.21 (-0.46-0) | Cagan et al. 2016 | 0-fold | MK test (McDonald and Kreitman 1991) |
|  |  | 0.000029 (0.0000032-0.01) | 0.0081 (0.0007-0.0911) |  | Castellano et al. 2019 | CDS | polyDFE v2.0 |
| Bonobo | Congo |  |  | -0.04 (-0.37-0.15) | Cagan et al. 2016 | 0-fold | MK test |
|  | Congo | 0.017 (0.000073-0.12) | 0.076 (0.0008, 27.8) |  | Castellano et al. 2019 | CDS | polyDFE v2.0 |
| Gorilla | Western lowland |  |  | 0.03 (-0.16-0.21) | Cagan et al. 2016 | 0-fold | MK test |
|  |  | 0.0013 (0.0000071-0.03) | 0.0091 (0.0008, 0.2291) |  | Castellano et al. 2019 | CDS | polyDFE v2.0 |
|  |  |  |  | 0.014 (-0.116-0.110) | McManus et al. 2015 | Nonsyn | DFE-alpha |
| Lemur | Coquerel's sifaka |  |  | 0.75 (0.7- 0.78) | Galtier 2016 | Nonsyn | Grapes |
| Mouse | Castaneus (mixed) |  |  | 0.32 (0.28-0.35) | Halligan et al. 2013 | 0-fold | DFE-alpha |
|  |  | 0.000179 (0.0000865-0.0002715) | 170.6 (89.7-251.55) |  | Booker et al. 2021 | CDS | Maximum likelihood |
|  |  | 0.003 (0.0019-0.0048) | 14.54 (9.24-23.4) |  | Booker et al. 2018 | 0-fold | DFE-alpha |
|  | Castaneus<br>(NW India) |  |  | 0.51 | Zhen et al. 2021 | Nonsyn | DFE-alpha |
|  |  | 0.04 | 3.76 | 0.41 (0.411-0.413) | Zhen et al. 2021 | Nonsyn | Composite likelihood approach |
| Rabbit | *Lepus granatensis* |  |  | 0.77 (0.68-0.86) | Galtier 2016 | Nonsyn | Grapes |
|  | *Oryctolagus cuniculus* |  |  | 0.64 (0.53-0.71) | Carneiro et al 2012 | Nonsyn | DFE-alpha |
| *Drosophila melanogaster* | Rwandan | 0.0045 (0-0.0165) | 23 (10-36) | 0.57 | Keightley et al. 2016 | Nonsyn | DFE-alpha |
|  | Yakuba | 0.00022; 0.000841<br>(high gene conversion rates) | 249; 508 (high gene conversion rates) |  | Campos et al. 2017 | Nonsyn | DFE-alpha |
|  | Zambian |  |  | 0.71 | Zhen et al. 2021 | Nonsyn | DFE-alpha |
|  |  | 0.04 | 4.92 | 0.60 (0.601-0.604) | Zhen et al. 2021 | Nonsyn | Composite likelihood approach |
|  |  |  |  | 0.59 | Lange & Pool 2018 | Nonsyn | asymptoticMK (Haller and Messer 2017) |
|  | East African | 0.00001 | 10000 |  | Jensen et al. 2008 | Nonsyn | ABC |
|  | Raleigh, NC |  |  | 0.40 | Elyashiv et al. 2016 | Nonsyn | Maximum composite likelihood |

## Results

### Neutrality with variable recombination/mutation rates in human-like populations

To assess the performance of *Relate* with realistic human data we performed detailed forward simulations of a whole chromosome, informed by empirical estimates, under five demographic histories: constant population size, recent expansion, recent contraction, bottleneck/recovery, and repeated bottleneck/recovery (Figure 1). Mutation and recombination rate heterogeneity was modelled across the chromosome according to published maps (The International HapMap Consortium 2005; Francioli et al. 2015) for all simulations (except where stated), rather than simulating a constant rate. Under equilibrium, (*i.e*., constant population size *N*), *Relate* inferred the correct demographic history and only slightly underestimated the population size up to 4*N* generations prior to sampling (Figure 1A). *Relate* effectively identified the presence of historical population expansion and bottleneck/recovery, however the precise timing and speed of the bottleneck and initial population growth/recovery was less accurate (Figure 1B, 1D). The presence, timing, and scale of population contraction was correctly identified under neutrality (Figure 1C), though a very recent brief increase in population size approximately 1000 generations from present was spuriously inferred prior to a sharp decrease toward the present (Figure S1); *i.e*., *Relate* correctly estimates the direction of population size change and the approximate scale of the overall decline in population size over *N*_*anc*_ (the ancestral population size) generations, though overestimates the rapidity of the decline in very recent generations for human-like populations. For the repeated bottleneck/recovery scenario *Relate* identified the presence of the most recent bottleneck/recovery, although the distant bottleneck/recovery was not inferred and elevated variation between replicates was seen along with mis-inference prior to the most recent bottleneck/recovery (Figure 1E). Our results suggest that mutation and recombination rate heterogeneity across the chromosome by itself had a negligible effect on simulated diversity and subsequent demographic inference for human simulations (Figure 1).

**Figure 1.**
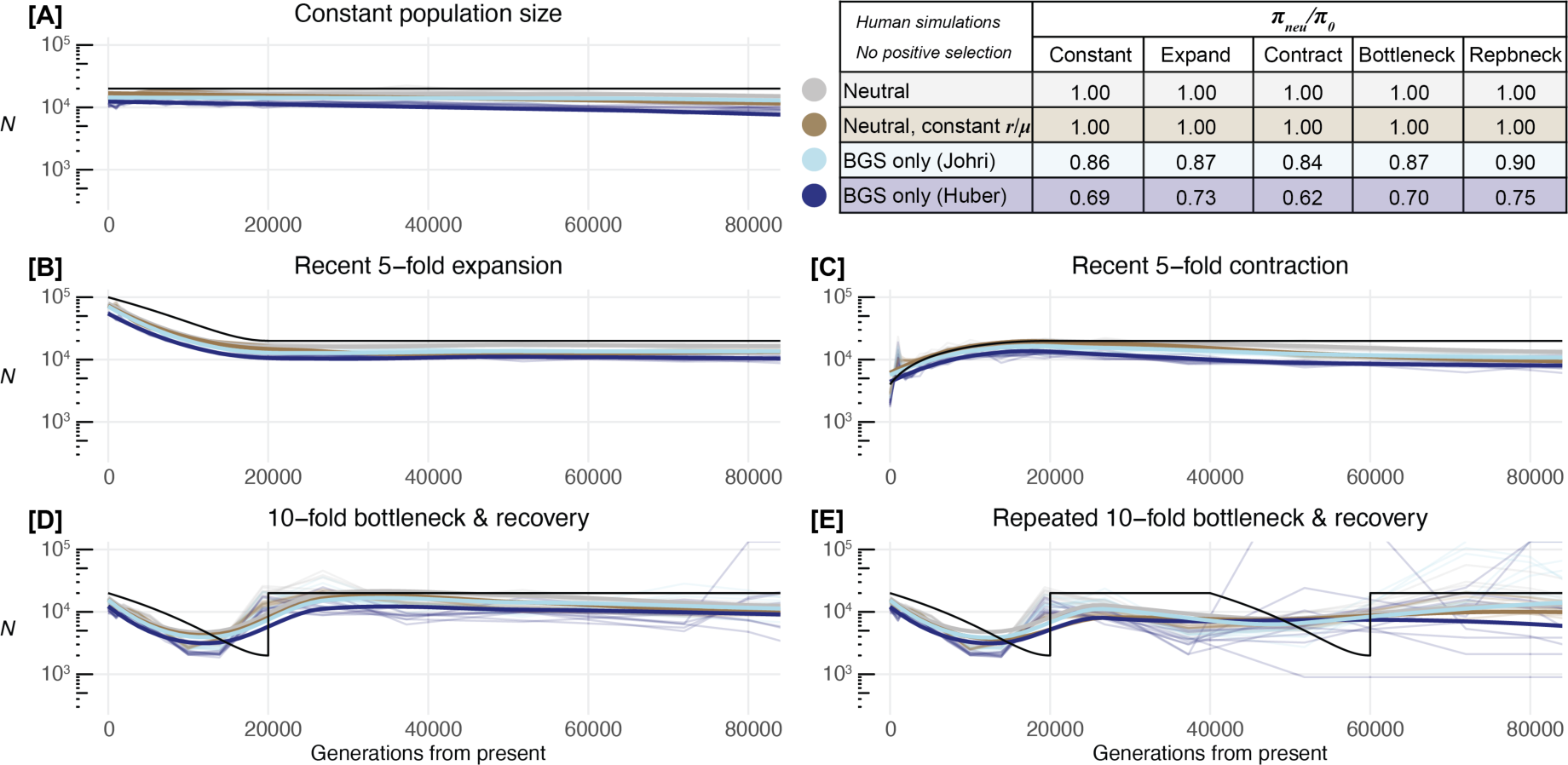
Historical population size inferred by *Relate* for human-like parameters under five demographic scenarios under neutrality and in the presence of BGS. The black line represents the true simulated population size (*N*) for each demographic scenario (A-E); coloured lines represent simulations without selection with varying recombination/mutation rates (grey), with constant recombination/mutation rates (gold), with BGS from the DFE reported by Johri *et al*. (2023; light blue) and with BGS from the DFE reported by Huber *et al*. (2017; dark blue). Thin coloured lines represent each of the 10 replicates per evolutionary scenario, thick coloured lines represent the moving regression (LOESS) across all replicates for a given condition. Nucleotide diversity at neutral intron/intergenic sites for simulations with selection relative to identical simulations under neutrality is presented as 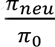 in the table. Note, population size is presented on a log_10_ scale.

### Background selection with variable recombination/mutation rates in human-like populations

BGS effects were modelled along with variable recombination and mutation rates. The simulated 138.6 Mb genome consisted of an exon-intron-intergenic structure, with the first and last exons of all genes flanked by UTR regions; realistic gene density heterogeneity was modelled by varying the distance between adjacent genes (see Methods). All coding and UTR regions experienced selection, while 4.4% and 3.6% of all new mutations in introns and intergenic regions respectively experienced selection, matching empirical estimates (Siepel et al. 2005; Dukler et al. 2022). This resulted in 3.15% of genome-wide sites experiencing selection in simulations of human-like populations, consistent with recent estimates of 3.2% of the human genome experiencing selection (Dukler et al. 2022). Purifying selection was modelled by simulating the distribution of fitness effects (DFE) estimated by Huber *et al*. (2017) and Johri *et al*. (2023), as discrete uniform distributions describing proportions of new mutations drawn from four classes of selective effects. While both estimates predict similar proportions of effectively neutral mutations (0 ≤ 2*N*_*anc*_*s*_*d*_ < 1), the DFE estimated by Johri *et al*. (2023) comprises more mild (1 ≤ 2*N*_*anc*_*s*_*d*_ ≤ 10) and moderately (10 ≤ 2*N*_*anc*_*s*_*d*_ < 100) deleterious mutations, and no strongly deleterious new mutations (2*N*_*anc*_*s*_*d*_ ≥ 100), compared to the Huber *et al*. (2017) DFE which predicts 22% of new mutations at conserved sites have strongly deleterious effects (see Methods).

The addition of purifying selection using the Huber *et al*. (2017) DFE reduced genetic diversity in neutral regions by 25-38% (Figure 1; Table S1, S2), with greater reductions for the population contraction scenario (38%) and lesser reductions for population expansion (25%). In comparison, we found the reduction of diversity under the Huber *et al*. (2017) DFE was greater than the 10-16% reduction under the DFE estimated by Johri *et al*. (2023; Figure 1; Table S1, S2). While the introduction of BGS led to an overall reduction in estimates of historical *N* inferred by *Relate* across all scenarios, as expected, the effect was relatively minor for human-like genomes and population parameters (Figure 1). While the true demographic history was largely recovered accurately in the presence of BGS in human-like populations, under the repeated bottleneck/recovery scenario there was considerable stochasticity between replicates in the more distant past as generations from present approached 4*N*_*anc*_ (Figure 1E).

### Effects of recurrent selective sweeps on demographic inference in human-like populations

Selective sweeps were modelled in addition to variable recombination and mutation rates and BGS effects by introducing beneficial mutations as an additional class in the DFE describing the 3.15% of chromosome-wide sites experiencing selection. To model the DFE of new beneficial mutations, we first performed a literature survey and collated previous estimates of relevant parameters from human populations (Table 1). While several studies have estimated the fraction of beneficial substitutions (*α*) in human populations, there are only two recent studies that have obtained estimates of the proportion of new mutations that are beneficial (*f*_*pos*_) and the distribution of selection coefficients of beneficial mutations (*s*_*a*_). However, these estimates vary widely because the two studies use very different data. Moreover, Zhen *et al*. (2020) demonstrated that a wide range of *f*_*pos*_ (proportion of new beneficial mutations at selected sites) and 2*N*_*anc*_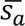 (mean population scaled selection coefficient of beneficial mutations) combinations could explain the observed substitution rates at nonsynonymous sites in humans across a likelihood surface. Thus, to explore a wide parameter space that could be relevant for human and great ape populations (see Table 1), we tested a range of *f*_*pos*_ and 2*N*_*anc*_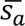 combinations in a grid-like fashion (Table 2). Beneficial mutations were introduced in simulations with one of two different proportions of new mutations (*f*_*pos*_= 0.01, *f*_*pos*_= 0.001); with each proportion, we tested population-scaled selection coefficients (2*N*_*anc*_*s*_*a*_) drawn from an exponential distribution with mean values of 10, 100, and 500. Note that our parameter combination *f*_*pos*_ = 0.01 and 2*N*_*anc*_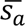 = 10 closely matches the empirical estimate provided by Zhen *et al*. (2020; Table 1).

The introduction of selective sweeps due to relatively rare beneficial mutations (*f*_*pos*_= 0.001), alongside BGS and variable recombination/mutation rates, resulted in effects on genetic diversity and demographic inference ranging from negligible to severe, depending on both the selection parameters and simulated demographic scenario (Figure 2; Table S1, S2). When the strength of positive selection was mild (2*N*_*anc*_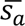 = 10, *f*_*pos*_= 0.001), negligible changes in nucleotide diversity and demographic inference were observed (Figure 2; Table S1, S2). Moderate, rare positive selection (2*N*_*anc*_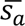 = 100, *f*_*pos*_= 0.001), resulted in moderate reductions in nucleotide diversity relative to strict neutrality (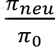 = 0.53-0.71; Figure 2; Table S1, S2), though demographic inference appeared largely unaffected.

**Figure 2.**
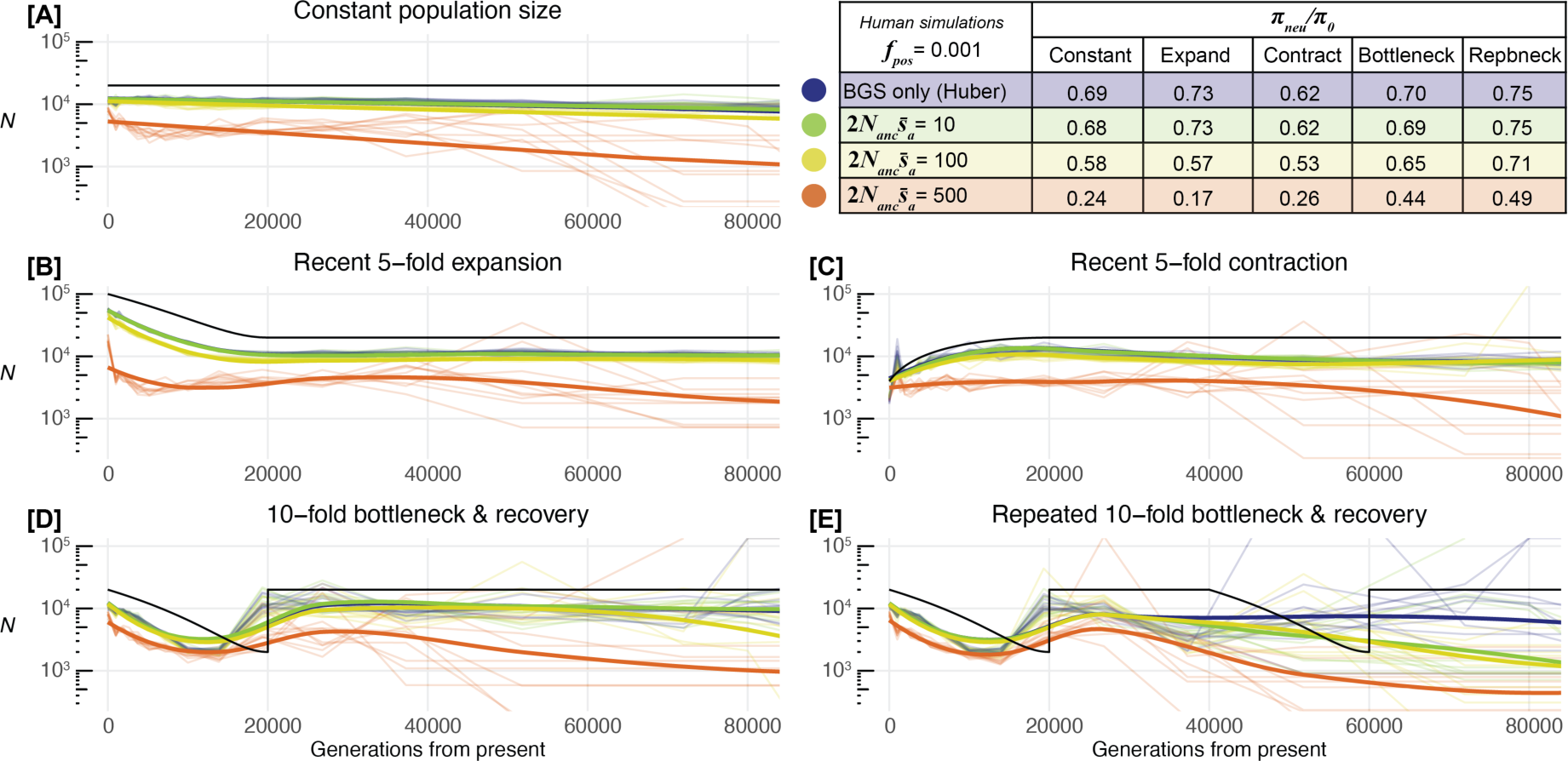
Historical population size inferred by *Relate* for human-like parameters under five demographic scenarios experiencing selective sweeps from relatively low frequency positive selection (*f*_*pos*_= 0.001) with variable recombination/mutation rates and BGS (DFE from Huber *et al*. 2017). The black line represents the true simulated population size (*N*) for each demographic scenario (A-E), coloured lines represent results for BGS only (blue), and different simulated mean positive selection coefficients (2*N*_*anc*_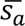). Thin coloured lines represent results for each of the 10 replicates per evolutionary scenario, thick coloured lines represent the moving regression (LOESS) across all replicates for a given condition. Nucleotide diversity at neutral intron/intergenic sites relative to identical simulations under neutrality is presented as 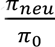 in the table. Note, population size is presented on a log_10_ scale.

Strong, rare positive selection (2*N*_*anc*_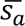 = 500, *f*_*pos*_= 0.001) resulted in large reductions in nucleotide diversity (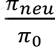 = 0.17-0.49; Figure 2; Table S1, S2) and led to demographic mis-inferences using *Relate*. For all demographic scenarios, strong, rare positive selection (2*N*_*anc*_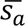 = 500; *f*_*pos*_= 0.001) resulted in gradual population expansion (approximately five-fold) from 4*N*_*anc*_ to around 1.5*N*_*anc*_ generations in the past, with increased stochasticity between replicates (Figure 2). In addition, it caused recent population expansion to be spuriously inferred under constant population size (Figure 2A), and constant population size to be incorrectly inferred under population contraction (Figure 2C). The presence of a recent population expansion was recovered under strong, rare positive selection (2*N*_*anc*_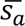 = 500, *f*_*pos*_= 0.001) as was a recent bottleneck, though large reductions in diversity and estimates of *N* were apparent, especially as generations from present approached 4*N*_*anc*_(Figure 2; Table S1, S2).

The introduction of selective sweeps due to relatively common beneficial mutations (*f*_*pos*_= 0.01), resulted in greater effects on genetic diversity and demographic inference than those produced under rarer occurrences of new beneficial mutations (*f*_*pos*_= 0.001). Mild, common positive selection (2*N*_*anc*_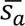= 10, *f*_*pos*_= 0.01) led to only slight reductions in neutral diversity (*π*_*neu*_) and did not impact demographic inference (Figure 3; Table S1, S2). Moderate strength, common positive selection (2*N*_*anc*_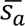 = 100, *f*_*pos*_= 0.01) led to large reductions in *π*_*neu*_ and led to poorly inferred demographic histories, including high stochasticity as generations from present approach 4*N*_*anc*_ and failure to recover a recent population contraction (Figure 3C), similar to effects seen for populations experiencing strong, rare positive selection (2*N*_*anc*_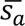 = 500, *f*_*pos*_= 0.001) parameter combination (Figure 2C). Strong, common positive selection (2*N*_*anc*_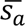 = 500, *f*_*pos*_= 0.01) resulted in extreme reductions of *π*_*neu*_ and failure to recover demographic histories, with severely underestimated *N* and spuriously inferred growth regardless of the simulated scenario (Figure 3). In summary, population history is mis-inferred by *Relate* when using neutral sites from human-like genomes if the combination of frequency (*f*_*pos*_) and strength of beneficial mutations (2*N*_*anc*_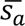) is sufficiently high (2*N*_*anc*_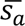*f*_*pos*_ ≥ 1).

**Figure 3.**
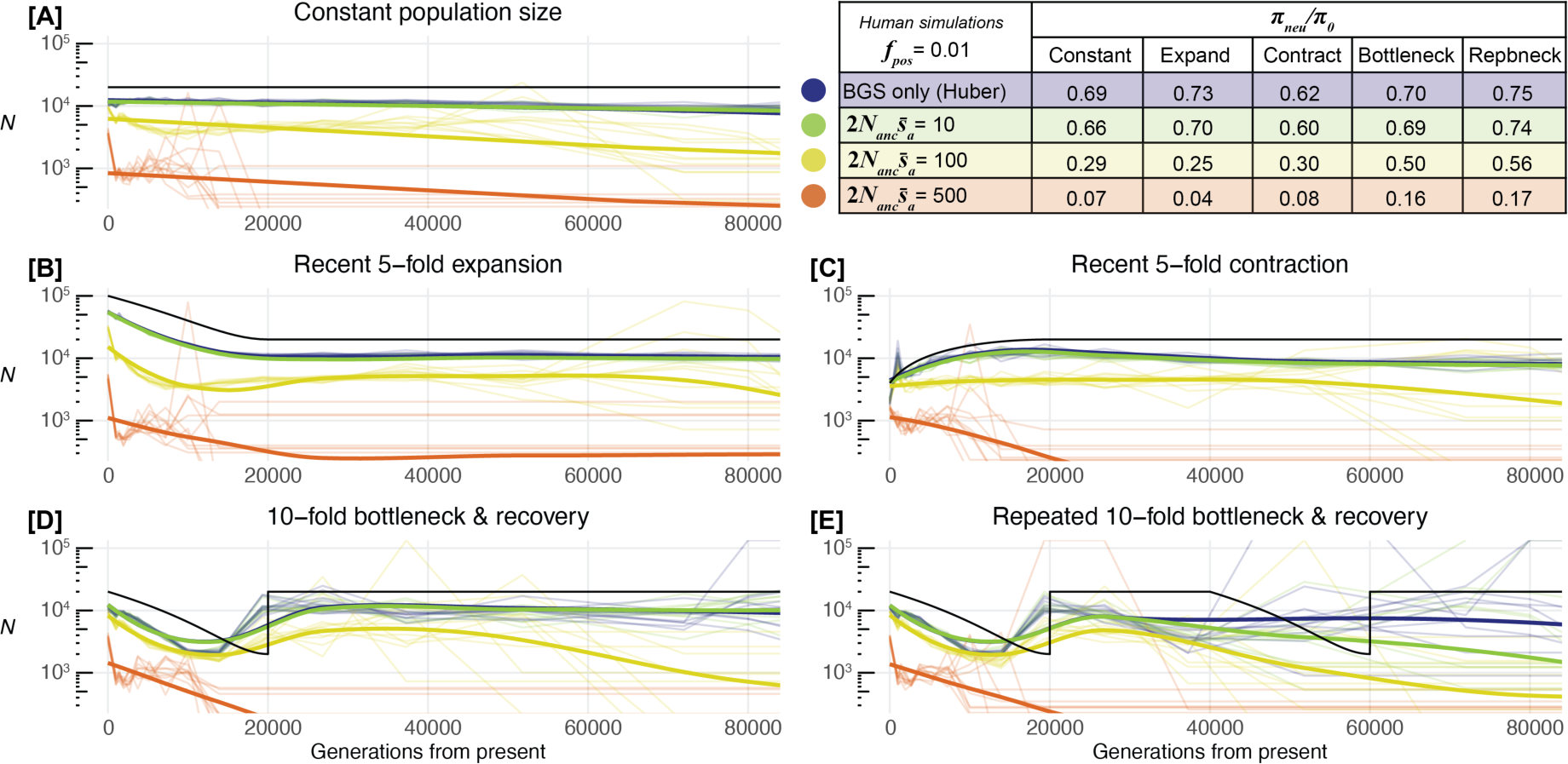
Historical population size inferred by *Relate* for human-like parameters under five demographic scenarios experiencing selective sweeps from relatively high frequency positive selection (*f*_*pos*_= 0.01) with variable recombination/mutation rates and BGS (DFE from Huber *et al*. 2017). The black line represents the true simulated population size (*N*) for each demographic scenario (A-E), coloured lines represent results for BGS only (blue), and different simulated mean positive selection coefficients (2*N*_*anc*_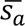). Thin coloured lines represent results for each of the 10 replicates per evolutionary scenario, thick coloured lines represent the moving regression (LOESS) across all replicates for a given condition. Nucleotide diversity at neutral intron/intergenic sites relative to identical simulations under neutrality is presented as 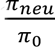 in the table. Note, population size is presented on a log_10_ scale.

### Relevance to human populations

While we have demonstrated that there exist parameter combinations where demographic inference will be biased in human populations in the presence of recurrent sweeps, it’s important to evaluate whether they are relevant to human populations. To assess the tenability of positive selection parameters for human simulations, we used the ratio of divergence at coding and neutral sites (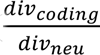), as used by (Soni et al. 2023), and the fraction of beneficial substitutions at coding sites, corrected to reflect values at nonsynonymous sites (*α*; Table 2; Table S3-S6), to compare our simulations with empirical findings (Table 1). In particular, our simulations indicated three combinations of parameters (2*N*_*anc*_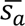 = 10, *f*_*pos*_= 0.001; 2*N*_*anc*_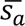 = 100, *f*_*pos*_= 0.001; 2*N*_*anc*_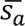= 10, *f*_*pos*_= 0.01) that result in *α* values within the published range for humans (0-0.25; Table 2) as well as plausible ratios of divergence between coding and neutral sites (Table 2). Thus, positive selection causing classic hard sweeps in humans is likely to be relatively rare and not strong, consistent with previous literature. Importantly, simulations with the parameter combinations that exhibited plausible summary statistics for human populations (Table 2) did not impact demographic inference with *Relate* (Figure 2, 3).

In contrast, when common positive selection was moderate or strong (2*N*_*anc*_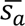 = 100, *f*_*pos*_= 0.01; 2*N*_*anc*_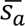 = 500, *f*_*pos*_= 0.01), parameter combinations that caused mis-inference with *Relate* (Figure 3), resulted in higher divergence in coding regions experiencing selection compared to neutral intergenic regions (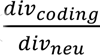> 1; Table 2), indicating that it is unlikely that these parameter combinations are relevant to human populations. Simulations with strong, rare positive selection (2*N*_*anc*_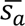 = 500, *f*_*pos*_= 0.001), which also caused mis-inference with *Relate* (Figure 2), exhibited plausible 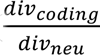 values (Table 2), with *α* values (0.40-0.47) exceeding published findings for humans and great apes, however this parameter combination may be relevant to rapidly evolving distant mammalian species with higher reported *α* values, such as mouse (Table 1).

### Effects of selection on demographic inference in *Drosophila melanogaster*-like genomes

To gauge the effects of BGS and selective sweeps on demographic inference with more compact genomes, we used similar approaches as described above for humans, with variable recombination/mutation rates, though simulated for *D. melanogaster*-like parameters (Table S7). The *D. melanogaster* genome architecture was simulated with 48.7% of genome-wide sites under selection, again distributed in exons, UTRs, introns (43% sites under selection), and intergenic (35% sites under selection) regions, consistent with estimates by Siepel *et al*. (2005). We found that providing an accurate mask input to *Relate* for recalculation of genetic distance was essential for accurate demographic inference for compact genomes where variants at a substantial proportion of total sites are removed from the analysis (48.7%; see Supplementary Note S1; Figure S2).

Introducing recombination rate and mutation rate heterogeneity led to a moderate improvement in demographic inference performance with *Relate* for *D. melanogaster* genomes across all demographic scenarios; *N* was relatively under-estimated by 2-fold to 10-fold in the distant past as generations from present approached 4*N*_*anc*_ for simulated populations with constant recombination and mutation rate when compared to simulated populations with realistic heterogeneous rates (Figure S3). For neutral simulations with realistic heterogeneity in mutation and recombination rates, *Relate* recovered the correct demographic histories under equilibrium (constant population size), population expansion, contraction, and the most recent bottleneck/recovery (Figure 4). However, approximately 2-fold variation in population size was inferred across the 4*N* period under equilibrium, as well as spurious inference of population expansion from ∼2500 generations ago until present with high stochasticity between replicates (Figure S4). In addition, an additional bottleneck further in the past was not recovered (Figure 4).

**Figure 4.**
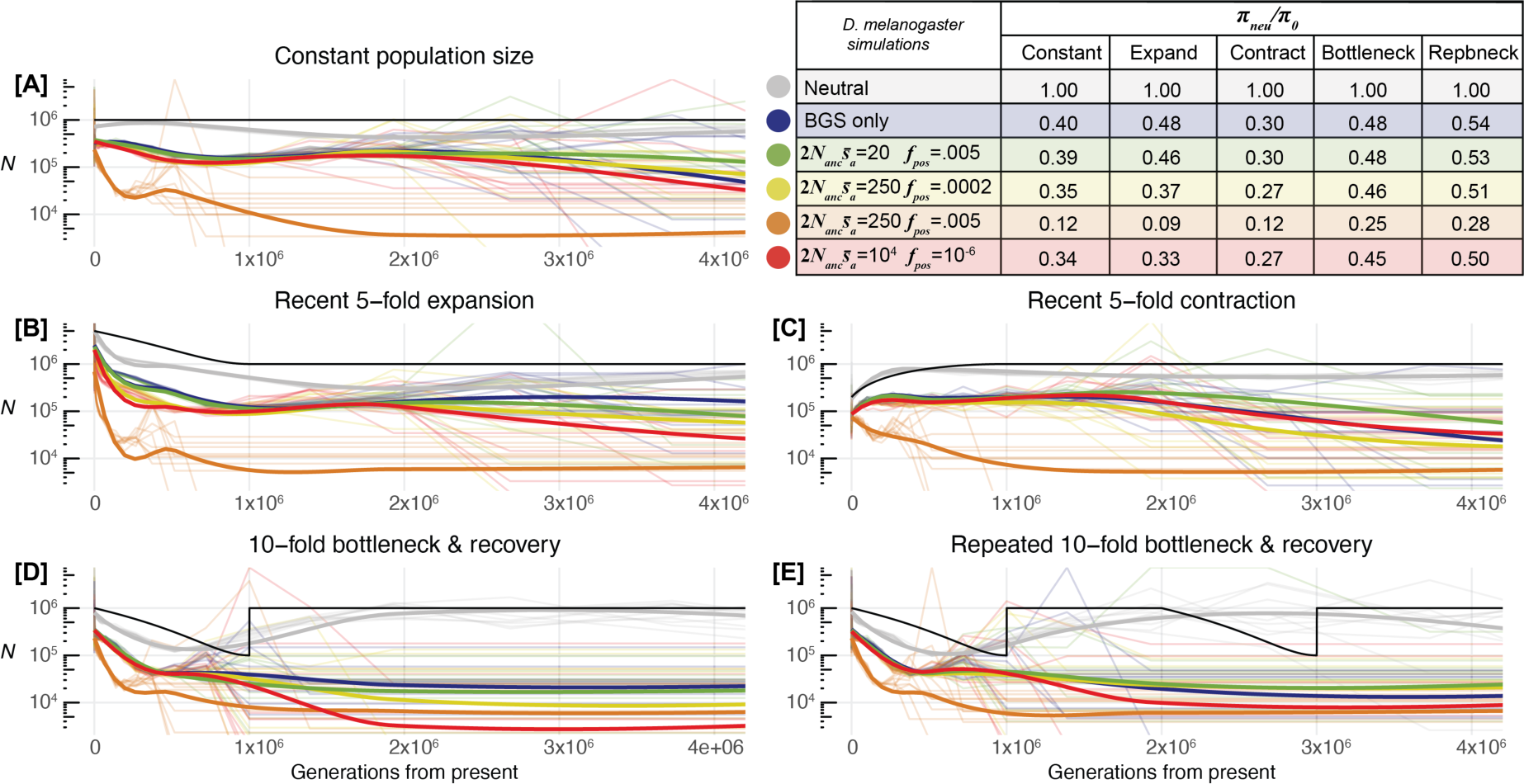
Historical population size inferred by *Relate* for *Drosophila melanogaster*-like parameters under five demographic scenarios, with variable recombination/mutation rates. The black line represents the true simulated population size (*N*) for each demographic scenario (A-E); coloured lines represent simulations under strict neutrality (grey), with BGS from the DFE reported by Johri *et al*. (2020; blue), and simulations experiencing BGS as well as sweeps from positive selection introduced with different parameter combinations. Thin coloured lines represent results for each of the 10 replicates per evolutionary scenario, thick coloured lines represent the moving regression (LOESS) across all replicates for a given condition. Nucleotide diversity at neutral intron/intergenic sites for simulations with selection relative to identical simulations under neutrality is presented as 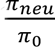 in the table. Note, population size is presented on a log_10_ scale.

Introducing BGS effects, modelled by applying the DFE estimated for deleterious mutations by Johri *et al*. (2020) to conserved sites (48.7% region-wide), reduced nucleotide diversity in neutral regions by 46-70% (Figure 4; Table S8, S9) consistent with theoretically predicted values by Charlesworth (2013). As expected, BGS led to substantially under-estimated *N* and increased stochasticity across replicates in all scenarios as generations from present approached 4*N*_*anc*_, with additional spurious variation in historical population size identified, even under equilibrium (Figure 4A). *Relate* recovered the recent population expansion and recent contraction, albeit with high stochasticity between replicates and underestimation of *N* beyond 1.5*N* generations ago for the population contraction scenario (Figure 4B, 4C). BGS caused recent bottlenecks to be entirely mis-inferred as recent expansions with population sizes under-estimated by up to two orders of magnitude prior to the bottleneck (Figure 4D, 4E).

Selective sweeps were modelled by introducing beneficial mutations at conserved sites with varying frequency (*f*_*pos*_) and mean positive selection coefficient (2*N*_*anc*_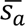), informed by widely varying empirically estimated values (Table 1), with BGS present. In contrast to human populations, several previous studies have estimated the relevant parameters in *D. melanogaster* populations (Table 1) though estimates vary widely, possibly due to methodological differences. We therefore simulated all estimated parameter combinations. The parameter combination (*f*_*pos*_= 0.005, 2*N*_*anc*_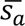 = 20) simulating common, weakly beneficial mutations reported by Keightley *et al*. (2016) had minimal additional effects on reducing diversity and impacting demographic inference, when compared to simulations with BGS only (Figure 4). The Campos *et al*. (2017) parameter combination (*f*_*pos*_= 0.0002, 2*N*_*anc*_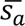 = 250) simulating infrequent, moderately beneficial mutations had a larger effect on reducing diversity, particularly in the population expansion scenario (Figure 4B), though demographic inference was only minorly impacted by the addition of sweeps. The Jensen *et al*. (2008) parameters (*f*_*pos*_= 1 × 10^−6^, 2*N*_*anc*_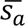 = 10000) simulating very rare, very strongly beneficial mutations had only moderate effects on reducing diversity and impacting demographic inference, though stochasticity between replicates was considerably higher in all scenarios, particularly when a recent population bottleneck occurred 1*N* generations ago (Figure 1D, 1E). However, simulated data from all three empirically estimated combinations of positive selection parameters produced summary statistics that did not match well with empirical findings: simulated 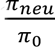 ranges from 0.54-0.27 (Figure 4, Table S8, S9), while Elyashiv et al. (2016) estimated 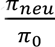 = 0.11-0.23; simulated *α* ≤ 0.19 (Table 2; Table S10) while published estimates report *α* = 0.40-0.71 (Table 1); and finally, while the simulated 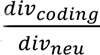 ≤ 0.52 (Table 2, S11-13), which is plausible in real populations experiencing selection, Johri *et al*. (2020) empirically estimated a higher ratio for *D. melanogaster* (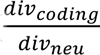= 0.87).

**Table 2.** Proportion of positively selected “nonsynonymous” substitutions at coding sites, corrected to reflect values at nonsynonymous sites (*α*) and the ratio of divergence in coding and neutral regions 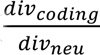 for human and *D. melanogaster* simulations experiencing different combinations of positive selection parameters: proportion of advantageous (*f*_*pos*_) and mean population scaled selection coefficient of advantageous mutations (2*N*_*anc*_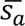) following an exponential distribution. Minimum and maximum values are shown, displaying the range across the five demographic scenarios tested for each parameter combination (Table S3-S6). Cells shaded green indicate simulated parameter combinations exhibiting higher divergence in neutral compared to coding regions, as expected in real populations, and *α* consistent with empirical observations in human and *D. melanogaster* populations (Table 1).

| Human simulations | | $2N_{anc}\bar{s}_a$ | | |
| --- | --- | --- | --- | --- |
|  |  | 10 | 100 | 500 |
| $f_{pos}$ | 0.001 | $\frac{div_{coding}}{div_{neu}} = 0.54-0.55$ | $\frac{div_{coding}}{div_{neu}} = 0.59-0.63$ | $\frac{div_{coding}}{div_{neu}} = 0.80-0.89$ |
| | | $\alpha = 0.02-0.03$ | $\alpha = 0.16-0.24$ | $\alpha = 0.40-0.47$ |
| | 0.01 | $\frac{div_{coding}}{div_{neu}} = 0.59-0.61$ | $\frac{div_{coding}}{div_{neu}} = 0.99-1.13$ | $\frac{div_{coding}}{div_{neu}} = 1.58-1.70$ |
| | | $\alpha = 0.16-0.23$ | $\alpha = 0.58-0.65$ | $\alpha = 0.71-0.74$ |
| *D. melanogaster* simulations | | $2N_{anc}\bar{s}_a$ | | |
|  |  | 20 | 250 | 10000 |
| $f_{pos}$ | 0.005 | $\frac{div_{coding}}{div_{neu}} = 0.48-0.52$ | $\frac{div_{coding}}{div_{neu}} = 0.91-0.98$ | |
| | | $\alpha = 0.11-0.19$ | $\alpha = 0.40-0.47$ | |
| | 0.0002 | $\frac{div_{coding}}{div_{neu}} = 0.43-0.49$ | $\frac{div_{coding}}{div_{neu}} = 0.48-0.52$ | |
| | | $\alpha = 0.01$ | $\alpha = 0.06-0.11$ | |
| | 0.000001 | | | $\frac{div_{coding}}{div_{neu}} = 0.46-0.51$ |
| | | | | $\alpha = 0.01$ |

The parameter combination of *f*_*pos*_ = 0.005 (Keightley et al. 2016) and 2*N*_*anc*_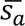= 250 (Campos et al. 2017) simulating common, moderately beneficial mutations, which was never inferred by any one previous study, produced summary statistics (*α*, 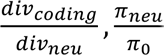; Table S8-S13) that best matched empirical estimates, which could potentially suggest that it most accurately models the effects of selection in real *D. melanogaster* populations. This parameter combination resulted in total reduction of neutral diversity (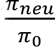 = 0.09-0.28; Figure 4, Table S8, S9) consistent with findings by Elyashiv *et al*. (2016), as well as *α* (0.40-0.47; Table 2, S10) and 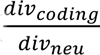(0.91-0.98; Table 1, S11-S13) best matching empirical estimates (Keightley et al. 2016; Campos et al. 2017; Lange and Pool 2018; Johri et al. 2020; Zhen et al. 2021) of all simulated *D. melanogaster* parameter combinations. Selective sweeps introduced by this parameter combination (*f*_*pos*_= 0.005, 2*N*_*anc*_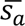= 250), with BGS present, led to failure by *Relate* to effectively infer demographic histories, with recent population expansion spuriously inferred in all simulated scenarios, severe underestimates of *N*, and extremely high stochasticity between replicates, even under equilibrium (Figure 4).

## Discussion

### Implications for human-like populations

Our analysis of realistic simulated populations (not accounting for structure) indicated that signatures of the linked effects of selection have the potential to cause mis-inference of historical population size using *Relate* when purifying and/or positive selection are sufficiently strong and pervasive. However, demographic inference with empirically supported parameters for human populations was shown to not be strongly impacted by selection under most tested demographic models. While BGS introduced by the DFEs estimated by Huber et al. (2017) and Johri et al. (2023) had moderate effects reducing genomic variation for simulated human populations (*B* ≈ 0.7-0.9; Table S1, S2), consistent with previous estimates (*B* = 0.83; Murphy et al. 2023), our findings suggest the effect of BGS on demographic inference for humans is minimal, inducing only slight under-estimation of historical population size without causing spurious inferences. Selective sweeps from recurrent positive selection similarly had little effect on demographic inference with *Relate* when restricted to empirically supported parameters for human populations (Eyre-Walker and Keightley 2009; Galtier 2016; Uricchio et al. 2019; Zhen et al. 2021), only potentially impacting the inferred history for populations that experienced repeated bottlenecks.

For simulated populations experiencing recurrent selective sweeps with parameters (2*N*_*anc*_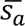, *f*_*pos*_, *α*) consistent with findings for human and other great apes with similar selection (McManus et al. 2015; Cagan et al. 2016; Castellano et al. 2019) and population-genetic dynamics (Besenbacher et al. 2019; Schmidt et al. 2019), we found the greatest driver of population history mis-inference using *Relate* was not selection but rather repeated fluctuations in historical population size. We found that when a simulated population experienced repeated bottlenecks *Relate* was only able to detect the most recent bottleneck, and exhibited poor inference further in the past, raising concerns of its accuracy when applied to many real populations. Great ape taxa including orangutans, gorillas, chimpanzees, and bonobos have been reported by some to experience a recent bottleneck (McManus et al. 2015) and repeated population bottlenecks (Prado-Martinez et al. 2013), suggesting that perhaps other demographic inference methods may be better suited at recovering repeated bottlenecks. Previous archaeological evidence suggests that we expect natural human populations underwent multiple bottlenecks (reviewed in Hawks et al. 2000; Nielsen et al. 2017). Our results suggest that while recent bottlenecks humans experienced may be observable (*e.g.,* Gravel et al. 2011; Liu and Fu 2015; Terhorst et al. 2017; Almarri et al. 2021; Ragsdale et al. 2023), *Relate* and similar methods may not effectively identify additional distant bottlenecks that may have taken place (Hawks et al. 2000).

While recurrent selective sweeps using parameters supported by findings for humans (2*N*_*anc*_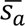 = 10, *f*_*pos*_= 0.001; 2*N*_*anc*_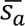 = 100, *f*_*pos*_= 0.001; 2*N*_*anc*_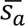 = 10, *f*_*pos*_= 0.01) are not strong and pervasive enough to meaningfully impact demographic inference with *Relate*, we found that stronger positive selection (2*N*_*anc*_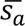 = 500, *f*_*pos*_= 0.001) could lead to mis-inference, which may be relevant to other mammalian species. Lemur (*Propithecus coquereli*; *α* = 0.75; Galtier 2016), mouse (*Mus musculus; α* = 0.41, 0.51; Halligan et al. 2013; Zhen et al. 2021), and rabbit populations (*Lepus granatensis, Oryctolagus cuniculus;* Carneiro et al. 2012; Galtier 2016) have been estimated to have a high fraction of beneficial relative to neutral substitutions at or above levels which caused *Relate* to produce spurious population expansions with substantial underestimations of historical population size (*α* = 0.40-0.47; Figure 4, Table S13). Mice and rabbits have higher gene densities than humans (Carneiro et al. 2014; Frankish et al. 2019) and larger effective population sizes (Carneiro et al. 2011; Phifer-Rixey et al. 2012), which would exacerbate the impact of both BGS and sweeps on demographic inference, though they also exhibit higher rates of recombination relative to mutation rates (Jensen-Seaman et al. 2004; Booker et al. 2017; Lindsay et al. 2019). Overall, our results indicate that selection has the potential to cause mis-inference of population history with *Relate* in some mammalian populations, though additional simulations modelling alternative genomes/populations are needed to identify the precise effects for non-human species. In cases where ARG-based inference using all intergenic/intronic sites are expected to be impacted due to the linked effects of selection, masking only selected regions may be insufficient and instead it may also be necessary to mask adjacent regions most affected by BGS and selective sweeps.

### Implications for *Drosophila*-like species with compact genomes

Our results suggest that demographic inference with *Relate* is susceptible to mis-inference for species with compact genomes and pervasive selection, as modelled by our *D. melanogaster* simulations. BGS due to purifying selection introduced according to the DFE estimated by Johri et al. (2020) substantially impacted demographic inference and decreased historical population size estimates by approximately ten-fold 1*N*_*anc*_ generations ago. Interestingly, the recent approximately two-fold expansion in the most recent 1*N*_*anc*_generations inferred by *Relate* for a population of constant size experiencing BGS (Figure 4A) matches well empirical estimates of two-to four-fold growth in the Zambian population of *D. melanogaster* (Ragsdale and Gutenkunst 2017; Kapopoulou et al. 2018), raising questions of whether these findings of population growth may be equally explained by a population of constant size experiencing the linked effects of selection. In addition, the effects of BGS in *D. melanogaster* resulted in the failure of *Relate* to recover some demographic shifts including population bottlenecks. Various non-Zambian *D. melanogaster* populations experienced severe bottlenecks, such as during transcontinental colonization events (Thornton and Andolfatto 2006; Laurent et al. 2011; Duchen et al. 2013; Kapopoulou et al. 2018; Arguello et al. 2019). Our results indicate that BGS alone is sufficient to entirely obscure the presence of population bottleneck when inferred by *Relate*, instead spuriously inferring a recent population expansion.

Selective sweeps will likely also exacerbate mis-inference in *D. melanogaster*, however, the extent to which this occurs may be unclear, as our simulated parameters matching empirical estimates varied widely in their frequency/intensity of selection and their effect on demographic inference. The mismatch we found between summary statistics (*α*, 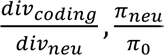; Table S8-S13) from our simulations using widely varying published *f_pos_* and 2*N*_*anc*_*s*_*a*_combinations (Jensen et al. 2008; Keightley et al. 2016; Campos et al. 2017; Zhen et al. 2021) compared to the same summary statistics reported in the literature (Elyashiv et al. 2016; Keightley et al. 2016; Campos et al. 2017; Johri et al. 2020; Zhen et al. 2021) raises concerns of consensus regarding which parameters best describe real populations. This is likely in part due to the likelihood surface as presented in Zhen *et al*. (2021), which highlighted the difficulty in accurately inferring the fraction of new beneficial mutations and their strength jointly. Another possibility is that the beneficial DFE in real *Drosophila* populations may deviate from the expected exponential distribution (Fisher 1930; Gillespie 1984; Orr 2003) modelled in our simulations and inferred by previous studies, and may instead be better fit by alternative distribution domains (Joyce et al. 2008; Seetharaman and Jain 2014). Similarly, while we assumed that all new mutations are semidominant, increased recessivity or dominance of selected mutations could result in different effects on genomic variation (Teshima and Przeworski 2006). Thirdly, it is possible that the large rescaling of *D. melanogaster*-like simulated populations that was necessary for computational tractability (see Methods; Table S14) inadvertently changed some dynamics of recombination by under-modelling crossover interference which increases the probability of introducing more than one crossover events in a single progeny; in such a case, increased linkage effects between distal sites may occur which could in turn influence selection dynamics. Generally, increased linkage and a higher mutation rate per diploid genome is likely to cause interference between selected alleles, leading to a possible decrease in the efficacy of positive selection.

Alternatively, it is possible that diminished effects of recombination could lead to increased effects of background selection, resulting in a lower effective population size, and therefore reduced efficacy of positive selection (*i.e*., lower values of *N*_*e*_*s*_*a*_). While we optimized our simulations based on the best fit to empirical findings within the realms of tractability, the precise impacts of rescaling are not fully known and further research is needed to clarify its effects and how best to preserve expected population genetics dynamics. Finally, positive selection in *D. melanogaster* and other populations may be primarily due to polygenic selection or selection on standing variation, in which case our assumed model of recurrent classic sweeps might not be sufficient to fit the empirical observations. Because positive selection on new mutations causes the strongest hitchhiking effect, it is unlikely that other models of positive selection will result in more severe mis-inference than that simulated in our study.

ARG-based demographic inference methods including *Relate* were initially developed to investigate human ancestry, however, these approaches have recently been applied to other organisms including insects, nematodes, and plants. Pope *et al*. (2023) reported population histories using ARG information approximated by *Relate*, inferring recent expansions in squash bee (*Eucera pruinosa*) populations following population splits coinciding with historical events. Our *D. melanogaster* results under the population bottleneck scenario (Figure 4D) suggest mis-inference due to BGS effects could partially explain some of their findings, if their methods for masking sites putatively linked to selected loci were too permissive (Pope et al. 2023; see Supplementary Note S2 for details). Beyond insects, *Relate* has recently been used to infer the demographic histories of self-fertilizing (selfing) and outcrossing *Caenorhabditis* nematodes (Teterina et al. 2023), and a grass species (*Brachypodium distachyon*) with high rates of selfing (Minadakis et al. 2023). Selfing increases the effects of background selection (Charlesworth et al. 1993; Charlesworth 2003) and selective sweeps (Hedrick 1980; Hartfield and Bataillon 2020), dramatically affects heterozygosity (Charlesworth and Charlesworth 1987) and linkage (Wright et al. 2013; Noble et al. 2021), and ultimately, changes the genealogical patterns from which *Relate* derives its demographic estimates (Nordborg and Donnelly 1997; Nordborg 2000; Strütt et al. 2023). Our results underline the need for caution when applying methods, such as *Relate*, that were designed for and tested with human parameters, to species with differing genome architecture experiencing disparate population dynamics.

### Demographic inference with complex genomic architecture

While a moderate improvement in inference was seen by effectively modelling recombination and mutation rate variability across *D. melanogaster* genomes (Figure S3), simulating recombination and mutation rate heterogeneity did not have a large impact on demographic inference with *Relate* across human genomes, unlike observations in some previous studies (Sellinger et al. 2021; Boitard et al. 2022). Sellinger *et al*. (2021) found that local variation in mutation rate could lead to demographic mis-inference and spurious apparent population decline using SMC methods. Realistic variation in local mutation and recombination rates has been shown to impact demographic inference when using SFS-based methods (Soni et al. 2024). In addition, extreme rates of recombination (cM/Mb ≤ 0.01, cM/Mb ≥ 5) have been shown to result in mis-inference of historical population size as calculated from nucleotide diversity using *Relate* (Novo et al. 2022). However, Upadhya and Steinrücken (2022) suggested that demographic inference performance with *Relate* improves when recombination rates are variable due to the presence of low recombination rate regions from which exact local genealogies can be estimated, which could in part explain the moderate improvement provided by modelling mutation and recombination rate heterogeneity in *D. melanogaster* simulated populations (Figure S3). It is unclear whether recombination and mutation rate heterogeneity may have more substantial effects in different species, as recent reports suggest masking low-recombination regions in songbird species may improve demographic inference with *Relate* (Ishigohoka and Liedvogel 2024).

The feasibility of finding neutral sites for demographic inference, which have not been substantially affected by nearby linked sites under selection, is essential for accurate genetic inference. Messer and Petrov (2013) found a 5-fold recent expansion could be mis-inferred by DFE-*α* using synonymous sites under equilibrium due to BGS alone, with more severe mis-inference (8.8-fold expansion) when selective sweeps were also modelled, with parameters (2*N*_*anc*_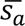 = 20, *α* = 0.18) similar to our empirically supported human parameters that did not lead to demographic mis-inference with *Relate* (Table 2; Figure 3A). Similarly, Schrider *et al*. (2016) revealed that when substantial fractions of neutral sites used for demographic inference are affected by proximal strong selective sweeps, population history can be severely impaired, concluding that it is essential to identify neutral sites sufficiently unlinked from recent selective sweeps to use for demographic inference. We found that neutral sites in intronic and intergenic regions are indeed sufficiently unlinked from recent selective sweeps to not affect inference in humans when simulating empirically supported positive selection parameters (Table 2; Figure 2, 3). However, our results suggest that the linked effects of selection (notably BGS) in compact genomes such as *D. melanogaster* impact demographic inference when using sites in neutral regions (Figure 4). Thus, for populations with compact genomes experiencing selection such as *D. melanogaster*, or in more extreme cases such as viruses where neutral sites unlinked to selected loci do not exist (Howell et al. 2023; Terbot et al. 2023), population history may be best inferred using joint inference approaches (*e.g.* Johri et al. 2020) that account for the linked effects of selection and demography simultaneously.

### The effect of selection on gene genealogies

Our results did not reveal that the effects of selection at linked sites lead to spurious inference of recent population decline, as reported by Boitard *et al*. (2022), when tested in realistic populations. The authors suggested the conflict between their findings (Boitard et al. 2022) and findings by Johri *et al*. (2021) was due to modelling of genomic regions without variable gene density or recombination and mutation rate variability; however, the simulated populations in our current analysis addresses these concerns and we largely corroborate findings by Johri *et al*. (2021). This is because Boitard *et al*. (2022) modelled the heterogeneity in effective population size across the genome while accounting for only the rescaling effects of *N*_*e*_ due to background selection and sweeps. However, both processes lead to not only a decrease in *N*_*e*_, but also affect the underlying gene genealogy, which mimics the genealogy of an expanding population and skews the SFS (Kim 2006; Nicolaisen and Desai 2013), and this is not accounted for by Boitard *et al*. (2022). Similarly, while Cousins *et al*. (2024) have recently implemented a *PSMC* method that accounts for the relative decrease in *N*_*e*_ across the genome caused by the linked effects of selection, such a method is unlikely to account for impacts of selection on local genealogies and thus may be ineffective when selection is pervasive, especially in species with compact genomes or those with little/no recombination. A final caveat of our analysis is the assumption of panmixia that may not reflect the reliability of ARG-based inference in the presence of population structure, which Boitard *et al*. (2020) showed could overshadow the impact of linked effects of selection on the IICR. Given reports that humans, for example, have experienced considerable population structure throughout their demographic history (Schiffels and Durbin 2014; Schlebusch et al. 2017; Ragsdale and Gravel 2019; Ragsdale et al. 2023), the effect of selection on inference in structured populations may be a promising avenue for additional research.

Following our analysis, the question arises: how generalizable are our findings using *Relate* to analysis with alternative ARG-based demographic inference approaches? Systematic ARG estimation using exhaustive likelihood approaches is computationally impractical (McVean and Cardin 2005), and as a result, *Relate* and similar methods use heuristics and HMMs to minimise redundant calculations (Li and Stephens 2003), greatly improving scalability with large sample sizes at the cost of precision. *Relate* (Speidel et al. 2019) and *tsinfer/tsdate* (Kelleher et al. 2019; Wohns et al. 2022) share biases when estimating recent and ancient coalescence times (Brandt et al. 2022; Upadhya and Steinrücken 2022), where accuracy drops when compared to *e.g*. the less scalable, more precise *ARGweaver* (Rasmussen et al. 2014), likely due to factors such as under-sampling of marginal trees (Deng et al. 2021). *Relate* has been shown in other studies to produce spurious inferences in very recent generation times, suggested to be due to a lack of sufficient mutations accumulating in short time intervals, as is needed for accurate inference using coalescent approaches, which alternate methods may improve upon (Santiago et al. 2020). Among the scalable approaches, *Relate* is relatively more affected by under-sampling of marginal trees (Deng et al. 2021), and preliminary reports indicated that demographic inference with *Relate* is less accurate than *tsinfer*/*tsdate* for the most recent ∼1000 generations, though *Relate* outperforms *tsinfer*/*tsdate* further in the past (Fan et al. 2023) while having improved coalescence time estimation than the recently developed *ARGneedle* (Ignatieva et al. 2023; Zhang et al. 2023). Thus, while minor differences in mis-inference due to effects of selection may occur across ARG-based methods, as the genealogy at linked neutral sites itself is altered by selection (Zeng and Charlesworth 2011; Nicolaisen and Desai 2013) we expect similar biases when performing demographic inference using branch lengths from alternative ARG approximations.

## Conclusions

ARG-based inference of population history remains a promising avenue for developing new methods (Brandt et al. 2024; Lewanski et al. 2024), however, *Relate* appears to be similarly impacted by the linked effects of selection as traditional SMC and SFS-based approaches (Johri et al. 2021). As a result, sufficiently pervasive positive and purifying selection leads to mis-inference of historical population size, most notably when moderately severe bottlenecks have taken place. While the effects of selection on demographic inference of human populations appears minimal, caution and additional testing with specific simulations are needed when analysing compact genomes with pervasive background selection and/or sufficiently high rates of positive selection, such as *D. melanogaster*, and populations with complex demographic histories.

Quantifying expected interactive effects between demography and selection on population variation remains necessary through the development of theoretical advances (Friedlander and Steinrücken 2022) and joint-inference approaches that account for the effects of selection at linked sites (Sheehan and Song 2016; Johri et al. 2020; Johri, Eyre-Walker, et al. 2022;). In addition, novel scalable ARG approximation with uncertainty estimates for coalescence times has recently been theorized (Wong et al. 2023), which would enable the development of improved methods that can account for temporal uncertainty (Brandt et al. 2024). While ARG-based demographic inference methods rapidly advance in terms of scalability and sophistication, it is essential we develop in tandem an understanding of their limits when applied to complex and varied empirical populations.

## Methods

### Simulation of chromosome structure

Simulations were performed using *SLiM* 4.0.1 (Haller and Messer 2023) with ten replicates for each evolutionary scenario. Simulations of human-like populations were performed assuming an ancestral population size (*N*_*anc*_) of 20,000 diploid individuals (Sherry et al. 1997; Huff et al. 2010), a mean mutation rate of 1.25 × 10^−/^(Kong et al. 2012), and a mean recombination rate of 1 × 10^−/^ per site/generation (Dumont and Payseur 2008; Kong et al. 2010). All *SLiM* simulations were scaled appropriately (population sizes and the number of generations were reduced, by a factor of 10 for human, while mutation and recombination rates were multiplied by the same factor) resulting in a scaled *N*_*anc*_ equal to 2000 (Comeron and Kreitman 2002; Hoggart et al. 2007; Kim and Wiehe 2008; Uricchio and Hernandez 2014). All simulations, human and *Drosophila*, included a burn-in period of 14*N*_*anc*_ generations, with an output of 50 diploid individuals (unless specified) in the VCF format. The number of output diploid individuals (50) used for demographic inference with *Relate* was chosen following testing with neutral simulations with human-like parameters which indicated that using 10 or fewer samples led to minor mis-inference in recent generations, while increasing the number of individuals beyond 50 up to 250 did not lead to substantial improvements in inference (Figure S5, S6). Note, we attempted to run *Relate* with 1000 samples, though results are not shown as demographic inference did not finish within 86 hours.

#### Drosophila

Simulations of *Drosophila melanogaster*-like populations were performed similarly, assuming *N*_*anc*_= 1 × 10^6^ (Keightley et al. 2014), a mean mutation rate of 3 × 10^−9^(Keightley et al. 2014), a mean recombination rate of 1 × 10^−8^ (Comeron et al. 2012). Given computational feasibility constraints of modelling *D. melanogaster* populations using forward-simulations with selection, we tested of a range of scaling factor and genome-size combinations (Table S14); as a result, we scaled simulations by a factor of 300 for a 5 Mb euchromatic region, which maximised the length of the region and resulted in empirically supported background selection effects (modelled as described below) in neutral regions: *B* = 0.4 simulated under constant population size (*B* = 0.45; estimated by Charlesworth 2013).

### Simulating the genome architecture

To approximate the genome architecture of the average human chromosome, we interspersed 797 genes with size and features approximately matching the mean values obtained across human protein-coding genes (Mignone et al. 2002; Mayr 2016; Piovesan et al. 2019). In simulated human genomes, all genes had 11 exons of 311 bp, 10 introns of 6938 bp, a 200 bp 5′ UTR, and an 800 bp 3′ UTR, with a mean intergenic length across the chromosome of 100 Kb. The 5′ UTR was placed immediately before the first exon in each gene; the 3′ UTR was placed immediately after the final exon in each gene; all genes had forward orientation.

To capture realistic variability of gene density, human gene features for chromosome 7 were downloaded from NCBI (https://www.ncbi.nlm.nih.gov; Sayers et al. 2023) using the following command: “*7[Chr] AND “Homo sapiens”[Organism] AND (“has ccds”[Properties] AND alive[prop])*”. The chromosome was divided into 1 Mb bins within which the number of genes were counted. The mean (5.47) and standard deviation (6.07) for the gene count across the 1 Mb bins were used to produce a normal distribution (limited to avoid negative gene counts and counts resulting in total genic lengths exceeding 1 Mb). We developed a script that recursively sampled 138 integers from this normal distribution (setSeed = 1), corresponding to the number of genes per 1 Mb bin, until the first occurrence of a cumulative sum (total gene count) of exactly 797 was output and saved to generate gene densities in simulations (see https://github.com/JohriLab/ARG_BGS_Demography/blob/main/scripts/gene_density_gener ation.slim). Within *SLiM*, the saved 138 integers were used to generate corresponding counts of gene structures in 1 Mb bins, by varying the intergenic lengths within each bin (Figure S7), with 300 Kb intergenic regions added to each end of the chromosome (*n.b.* the intergenic lengths are the only genomic elements that vary in length across the chromosome, and the chromosome structure was identical across all replicates and conditions). This produced a chromosome with a size of 138.6 Mb and genic regions (introns and exons) spanning 58 Mb (41.9%), which is consistent with the 1.27 Gb estimate of total protein coding gene length including introns (Piovesan et al. 2019), which spans 40.9% of a 3.12 Gb human genome (Nurk et al. 2022; Rhie et al. 2023).

#### Drosophila

The genome architecture of *Drosophila melanogaster* was simulated similarly, however, due to computational feasibility constraints, a 5 Mb euchromatic region was modelled based on features from chromosome arm 2R rather than a full chromosome. All 565 genes had 5 exons of 314 bp each, 4 introns of 999 bp each, a 266 bp 5′ UTR and a 382 bp 3′ UTR, with a mean intergenic length across the region of 2654 bp (Long and Deutsch 1999; Dos Santos et al. 2015; Table S7). The mean (14.69) and standard deviation (7.54) for gene count across 100 Kb bins was used to generate a normal distribution of gene counts in 50 variably dense 100 Kb bins totalling 565 genes over 5 Mb (see above and https://github.com/JohriLab/ARG_BGS_Demography/blob/main/scripts/gene_density_generation.slim), with 20 kb intergenic regions added to the ends of the region. This produced a 5.04 Mb region comprised of 17.70% coding sites, matching the chromosome 2R annotation for the BDGP6 assembly (17.68%; Dos Santos et al. 2015).

### Modelling heterogeneity in recombination and mutation rates

The sex-averaged mutation rate map for human chromosome 7 generated by Francioli *et al*. (2015) was used as input for the mutation rates, scaled so that the average mutation rate (*µ*) across the chromosome was 1.25 × 10^−8^ per site/generation (1.25 × 10^−1^ after scaling; see Figure S8 for distribution of mutation rates). As Francioli *et al*. (2015) provided the rates for each base pair transition (*i.e*., A to T, T to G, etc), the nucleotide composition for each corresponding 1 Mb interval was calculated from the GRCh37 reference (BioProject PRJNA31257) and multiplied by the sum of the rates for each given nucleotide. Finally, the rates for each nucleotide were combined to produce *µ* (see https://github.com/JohriLab/ARG_BGS_Demography/blob/main/scripts/mut_map.sh). The population-specific recombination rate map for chromosome 7 generated for the YRI population in the HapMap project (The International HapMap Consortium 2005) was used as input for the recombination rates. The shared length (146.04 Mb) reported in both the chromosome 7 recombination rate map and mutation rate map (7.01 Mb to 153.05 Mb) was extracted, and all interval lengths were reduced by approximately 5.1% to have a cumulative total length of 138.6 Mb rather than 146.04 Mb, so that the maximal region for which recombination rates and mutation rates are reported for chromosome 7 could be modelled in our simulations.

#### Drosophila

The mean mutation rate simulated was 3 × 10^−9^ per site/generation (Keightley et al. 2014) prior to scaling and was varied across the region in 50 kb bins. The mutation rates in each 50 kb bin were drawn from a normal distribution with a coefficient of variation of 0.479, corresponding to the coefficient of variation for divergence at 4-fold degenerate sites (*d*_*s*_) between *D. simulans* and *D. melanogaster* in the middle section of the 2R chromosome arm (Mackay et al. 2012). The mean sex-averaged recombination rate simulated (crossover probability per bp = 1 × 10^−8^) was modelled on a genome-wide estimate (Comeron et al. 2012), though rates were varied in 50 kb bins. The recombination rates in each 50 kb bin were taken from the rates in 100 Kb bins from 10 Mb to 20 Mb reported by Comeron *et al*. (2012) on the *Drosophila melanogaster* recombination rate calculator v2.30 (Fiston-Lavier et al. 2010).

### Simulating mutational fitness effects

Functional regions, coding sequences and UTRs, experienced purifying selection modelled by a distribution of fitness effects (DFE) comprised of four non-overlapping uniform distributions representing the effectively neutral (0 ≤ 2*N*_*anc*_*s*_*d*_ < 1), weakly deleterious (1 ≤ 2*N*_*anc*_*s*_*d*_ < 10), moderately deleterious (10 ≤ 2*N*_*anc*_*s*_*d*_ < 100), and strongly deleterious (100 ≤ 2*N*_*anc*_*s*_*d*_ < 2*N*_*anc*_) class of mutations, such that *f*_0_, *f*_1_, *f*_2_, and *f*_3_ proportion of all new mutations belonged to each class respectively. Here, *s*_*d*_ represents the selective disadvantage of mutant homozygotes relative to the wildtype, and all mutations were assumed to be semidominant. When simulating recurrent sweeps, an *f*_*pos*_ proportion of all new mutations in functional elements was assumed to be beneficial, while 1 − *f*_*pos*_ proportion was effectively neutral and deleterious. Beneficial mutations were assumed to be semidominant and to follow an exponential DFE with the mean selective advantage of mutant homozygotes relative to wildtypes denoted by 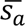. Semidominance of all mutations was assumed because previous estimates used to parameterize selection in our simulations were inferred under the same assumption.

The deleterious DFE estimated by Huber *et al*. (2017) for the Yoruba population was used to inform the proportions of each mutation class in human simulations where BGS was present: *f*_0_= 0.51; *f*_1_= 0.14; *f*_2_= 0.14; *f*_3_ = 0.21; the alternative Johri *et al*. (2023) DFE tested (Figure 1) was: *f*_0_ = 0.50; *f*_1_ = 0.20; *f*_2_ = 0.30; *f*_3_ = 0. Simulations that modelled recurrent selective sweeps included beneficial mutations with varying values of *f*_*pos*_, and the population-scaled selection coefficients of advantageous mutations (2*N*_*anc*_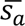), as provided in Table 2.

To model functional elements like regulatory elements present in intergenic and intronic regions, we assumed that conserved introns and intergenic regions experience selection. Previous findings report the functional proportion of the human genome experiencing purifying selection is approximately between 3.2%-8.2% (Siepel et al. 2005; Rands et al. 2014; Dukler et al. 2022). Approximately 60% of mutations with a deleterious fitness effect across the genome have been found to be located in intronic and intergenic regions (Siepel et al. 2005; Dukler et al. 2022). According to Table 2 in Dukler *et al*. (2022), 3.6% of intergenic regions, and 4.4% of intronic regions are conserved (i.e., containing elevated proportions of selected sites). Thus, instead of modelling the precise locations of regulatory elements, we assumed that 4.4% and 3.6% of all new mutations in intronic and intergenic regions were selected, following the same DFE as in coding and UTR regions, while the remaining majority of intergenic (96.4%) and intronic (95.6%) mutations were kept neutral (*n.b.* we did not explicitly model conserved non-coding element structures within introns and intergenic regions). Combined with the exon and UTR regions (2.54% of the genome) for which all new mutations possessed selection coefficients sampled from the DFE, the mean total proportion of new mutations across the genome with selection coefficients sampled from the DFE was 6.43%; given an effectively neutral (*f*_0_) proportion of 0.51, the conserved proportion of the genome ( *p_conserved_*) was 3.15%.

#### Drosophila

The structure of functional elements for *Drosophila* simulations was modelled as above, with constrained proportions of intron (43%) and intergenic (35%) sites, constrained proportion of all sites (48.7%) matching previous estimates (Andolfatto 2005; Siepel et al. 2005; Table S7). The deleterious DFE estimated by Johri *et al*. (2020) for the Zambian population was used for *Drosophila* simulations: *f*_0_ = 0.25; *f*_1_ = 0.49; *f*_2_ = 0.04; *f*_3_= 0.22. In total, 64.9% of new mutations across the genome sampled fitness effects from the DFE and *p_conserved_* was 48.7%, matching previous findings of 39%-53% (Siepel et al. 2005).

### Simulation of population history

Four demographic models were tested to assess the robustness of demographic inference results under a range of historical population scenarios following the burn-in period of 14*N*_*anc*_, at which time *N* = *N*_*anc*_ = 2000 for human simulations (3333 for *Drosophila*) after scaling. Recent expansion was modelled by exponential growth leading to a five-fold increase in population size that started 1*N*_*anc*_ generations ago. Recent contraction was modelled by exponential decline leading to a five-fold decrease in population size that started 1*N*_*anc*_ generations ago. Bottleneck and recovery were modelled by an instantaneous ten-fold decline in population size (1*N*_*anc*_ generations ago), which remained constant for 10 generations, before exponentially increasing to recover to *N*_*anc*_. Repeated bottlenecks were simulated identically to the bottleneck-and-recovery scenario, however following recovery, population size was kept constant for 1*N*_*anc*_ generations, before repeating the scenario.

### Calculating summary statistics

Mean nucleotide diversity in neutral regions (*π*_*neu*_) was calculated using the calcHeterozygosity function within *SLiM*, applied to only neutral mutations in the intergenic and intronic regions, multiplied by the total proportion of neutral sites in the genome. The decrease in nucleotide diversity in neutral regions due to the shared effects of background selection and selective sweeps under different selection parameters was estimated by 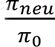 where *π*_0_ was obtained from otherwise identical simulations without selection.

Divergence at neutral sites in (*div*_*neu*_) was calculated as the number of neutral substitutions in intergenic and intronic regions that occurred after 10*N*_*anc*_ generations (*n.b.* all population size change scenarios took place after 14*N*_*anc*_burn-in) divided by the total number of neutral sites in intergenic and intronic regions (*i.e*., 95.65% of intronic and 96.40% of intergenic sites for human). Divergence in coding regions (*div*_*coding*_) was similarly calculated by dividing all exonic substitutions, neutral and selected, by the cumulative length of exon regions in the simulated chromosome.

The proportion of nonsynonymous substitutions in coding regions that were positively selected (*α*) was determined assuming ∼30% of mutations in coding regions are synonymous, and all synonymous mutations are neutral. Given ∼50% of new mutations in coding regions are neutral according to the human DFE provided by Huber *et al*. (2017), this suggests that the remaining ∼40% of neutral mutations in coding regions (∼20% of all exonic mutations) refer to non-synonymous sites. Therefore, *α* for simulated populations of humans was calculated by the number of substitutions observed in different classes of mutations (*sub*_*class*_) that occurred after 10*N*_*anc*_ using the following equation:

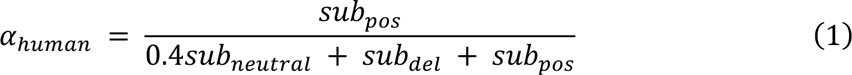

#### Drosophila

we assumed that all mutations in the effectively neutral class (comprising 25% of all mutations) following the Johri *et al*. (2020) DFE represent all synonymous mutations, and no nonsynonymous mutations are strictly neutral. Therefore, *α* for *Drosophila* was calculated as above, though without including any neutral substitutions:

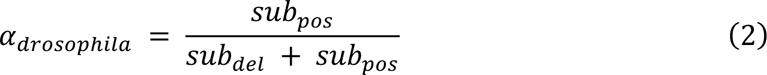

Scripts used to generate summary statistics are available at: https://github.com/JohriLab/ARG_BGS_Demography/blob/main/scripts/pi_div_alpha.sh.

### Demographic inference with *Relate*

The VCF for each replicate was filtered to only contain neutral mutations in intergenic and intronic regions, before being converted to the hap/sample format using the *RelateFileFormats* script, with *–mode ConvertFromVcf*. During testing, we found that when masking selected sites from the VCF used as input for *Relate*, *i.e.*, removing SNPs in selected regions, demographic inference could be majorly impacted if a mask file was not provided to *Relate* to enable recalculation of genetic distances between sites (see Supplementary Note S1 for details). As a result, we generated a genomic mask file for each individual replicate which masks conserved regions (exons/UTRs) as well as conserved sites and a randomly chosen proportion of invariant sites in introns/intergenic regions, to account for excluding/masking functional elements for demographic inference (see https://github.com/JohriLab/ARG_BGS_Demography/blob/main/scripts/new_build_mask.sh). The haps, sample, and mask files were then passed to *Relate* 1.1 (Speidel et al. 2019) to perform genome-wide genealogy estimation with mutation rate (*μ*) = 1.25 × 10^−8^for human (3 × 10^−9^for *Drosophila*). The estimated current haplotype population size (2*N*_*cur*_) parameter, ‘*-N*’, required by *Relate*, was calculated as may be done in empirical analysis, rather than using the true values from simulations, by the following:

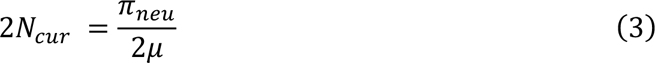

Where *μ* is the unscaled genome-wide mutation rate, and mean nucleotide diversity in neutral regions (*π*_*neu*_) is calculated as previously described. The genealogy results were passed through *Relate*’s EstimatePopulationSize module, with ‘--years_per_generation 1’. Historical effective population sizes for an epoch *i* generations ago (*N*_*i*_) were estimated from the coalescence rates output using the same method built into *Relate*:

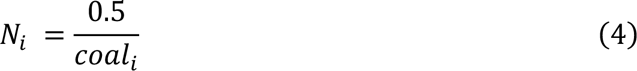

where *coal* is the mean genome-wide instantaneous coalescence rate *i* generations ago obtained from the *.coal* file output by *Relate*. The genetic maps were calculated from the recombination rate maps assuming linearity (see https://github.com/JohriLab/ARG_BGS_Demography/blob/main/scripts/rec_map2gen_map.sh); Haldane’s correction (Haldane 1919) was deemed unnecessary as mean interval length between rate recalculation < 1.1Mb*. n.b.* despite the *SLiM* simulations being scaled down, the *Relate* parameters are unscaled, thus producing unscaled demographic inference results.

Figure generation was performed using *R* v4.1.0 and *Python* v3.11.3. ChatGPT 3.5, (https://chat.openai.com; accessed September 2023) was used to build two minor data processing scripts (print_nums.sh and exon_mutations.py at https://github.com/JohriLab/ARG_Selection_Demography/blob/main/scripts/), though was not used for any text generation for the manuscript.

## Acknowledgements

The research in this study was conducted using computational resources provided by ITS Research Computing at the University of North Carolina at Chapel Hill. The authors declare no conflicts of interest. We thank Jeff Jensen for providing helpful comments on the manuscript.

## Data availability

All relevant code and scripts generated for this study are available at https://github.com/JohriLab/ARG_BGS_Demography. The mutation and recombination rate maps used for analysis are available at https://github.com/JohriLab/ARG_BGS_Demography/tree/main/misc.

## References

Almarri MA, Haber M, Lootah RA, Hallast P, Al Turki S, Martin HC, Xue Y, Tyler-Smith C. 2021. The genomic history of the Middle East. Cell 184:4612–4625.

Andolfatto P. 2005. Adaptive evolution of non-coding DNA in *Drosophila*. Nature 437:1149–1152.

Arguello JR, Laurent S, Clark AG. 2019. Demographic history of the human commensal *Drosophila melanogaster*. Genome Biol Evol 11:844–854.

Beaumont MA, Zhang W, Balding DJ. 2002. Approximate Bayesian computation in population genetics. Genetics 162:2025–2035.

Beichman AC, Huerta-Sanchez E, Lohmueller KE. 2018. Using genomic data to infer historic population dynamics of nonmodel organisms. Annu Rev Ecol Evol Syst 49:433–456.

Bergström A, McCarthy SA, Hui R, Almarri MA, Ayub Q, Danecek P, Chen Y, Felkel S, Hallast P, Kamm J, et al. 2020. Insights into human genetic variation and population history from 929 diverse genomes. Science 367:eaay5012.

Besenbacher S, Hvilsom C, Marques-Bonet T, Mailund T, Schierup MH. 2019. Direct estimation of mutations in great apes reconciles phylogenetic dating. Nat Ecol Evol 3:286–292.

Boitard S, Arredondo A, Chikhi L, Mazet O. 2022. Heterogeneity in effective size across the genome: effects on the inverse instantaneous coalescence rate (IICR) and implications for demographic inference under linked selection. Genetics 220:iyac008.

Booker TR, Jackson BC, Keightley PD. 2017. Detecting positive selection in the genome. BMC Biol 15:98.

Brandt D, Huber CD, Chiang CWK, Ortega-Del Vecchyo D. 2024. The promise of inferring the past using the ancestral recombination graph. Genome Biol Evol 16:evae005.

Brandt D, Wei X, Deng Y, Vaughn AH, Nielsen R. 2022. Evaluation of methods for estimating coalescence times using ancestral recombination graphs. Genetics 221:iyac044.

Cagan A, Theunert C, Laayouni H, Santpere G, Pybus M, Casals F, Prüfer K, Navarro A, Marques-Bonet T, Bertranpetit J, et al. 2016. Natural selection in the great apes. Mol Biol Evol 33:3268–3283.

Caicedo AL, Williamson SH, Hernandez RD, Boyko A, Fledel-Alon A, York TL, Polato NR, Olsen KM, Nielsen R, McCouch SR, et al. 2007. Genome-wide patterns of nucleotide polymorphism in domesticated rice. PLoS Genet 3:e163.

Campos JL, Zhao L, Charlesworth B. 2017. Estimating the parameters of background selection and selective sweeps in *Drosophila* in the presence of gene conversion. Proc Natl Acad Sci USA 114:E4762–E4771.

Carneiro M, Afonso S, Geraldes A, Garreau H, Bolet G, Boucher S, Tircazes A, Queney G, Nachman MW, Ferrand N. 2011. The genetic structure of domestic rabbits. Mol Biol Evol 28:1801–1816.

Carneiro M, Albert FW, Melo-Ferreira J, Galtier N, Gayral P, Blanco-Aguiar JA, Villafuerte R, Nachman MW, Ferrand N. 2012. Evidence for widespread positive and purifying selection across the European rabbit (*Oryctolagus cuniculus*) genome. Mol Biol Evol 29:1837–1849.

Carneiro M, Rubin C-J, Di Palma F, Albert FW, Alföldi J, Barrio AM, Pielberg G, Rafati N, Sayyab S, Turner-Maier J, et al. 2014. Rabbit genome analysis reveals a polygenic basis for phenotypic change during domestication. Science 345:1074–1079.

Castellano D, Macià MC, Tataru P, Bataillon T, Munch K. 2019. Comparison of the full distribution of fitness effects of new amino acid mutations across great apes. Genetics 213:953–966.

Charlesworth B. 2013. Background selection 20 years on. J Hered 104:161–171.

Charlesworth B, Morgan MT, Charlesworth D. 1993. The effect of deleterious mutations on neutral molecular variation. Genetics 134:1289–1303.

Charlesworth D. 2003. Effects of inbreeding on the genetic diversity of populations. Philos Trans R Soc B 358:1051–1070.

Charlesworth D, Charlesworth B. 1987. Inbreeding depression and its evolutionary consequences. Annu Rev Ecol Syst 18:237–268.

Chikhi L, Rodríguez W, Grusea S, Santos P, Boitard S, Mazet O. 2018. The IICR (inverse instantaneous coalescence rate) as a summary of genomic diversity: insights into demographic inference and model choice. Heredity 120:13–24.

Comeron JM, Kreitman M. 2002. Population, evolutionary and genomic consequences of interference selection. Genetics 161:389–410.

Comeron JM, Ratnappan R, Bailin S. 2012. The many landscapes of recombination in *Drosophila melanogaster*. PLoS Genet 8:e1002905.

Cousins T, Tabin D, Patterson N, Reich D, Durvasula A. 2024. Accurate inference of population history in the presence of background selection. Available from: http://biorxiv.org/lookup/doi/10.1101/2024.01.18.576291

Deng Y, Song YS, Nielsen R. 2021. The distribution of waiting distances in ancestral recombination graphs. Theor Pop Biol 141:34–43.

Duchen P, Živković D, Hutter S, Stephan W, Laurent S. 2013. Demographic inference reveals African and European admixture in the North American *Drosophila melanogaster* population. Genetics 193:291–301.

Dukler N, Mughal MR, Ramani R, Huang Y-F, Siepel A. 2022. Extreme purifying selection against point mutations in the human genome. Nat Commun 13:4312.

Dumont BL, Payseur BA. 2008. Evolution of the genomic rate of recombination in mammals. Evolution 62:276–294.

Elyashiv E, Sattath S, Hu TT, Strutsovsky A, McVicker G, Andolfatto P, Coop G, Sella G. 2016. A genomic map of the effects of linked selection in *Drosophila*. PLoS Genet 12:e1006130.

Ewing GB, Jensen JD. 2016. The consequences of not accounting for background selection in demographic inference. Mol Ecol 25:135–141.

Excoffier L, Marchi N, Marques DA, Matthey-Doret R, Gouy A, Sousa VC. 2021. *fastsimcoal2* : demographic inference under complex evolutionary scenarios. Bioinformatics 37:4882–4885.

Eyre-Walker A, Keightley PD. 2009. Estimating the rate of adaptive molecular evolution in the presence of slightly deleterious mutations and population size change. Mol Biol Evol 26:2097–2108.

Fan C, Cahoon JL, Dinh BL, Ortega-Del Vecchyo D, Huber C, Edge MD, Mancuso N, Chiang CWK. 2023. A likelihood-based framework for demographic inference from genealogical trees. Available from: http://biorxiv.org/lookup/doi/10.1101/2023.10.10.561787

Fisher RA. 1930. The genetical theory of natural selection. Oxford, UK: Clarendon Press

Fiston-Lavier A-S, Singh ND, Lipatov M, Petrov DA. 2010. *Drosophila melanogaster* recombination rate calculator. Gene 463:18–20.

Francioli LC, Polak PP, Koren A, Menelaou A, Chun S, Renkens I, Van Duijn CM, Swertz M, Wijmenga C, Van Ommen G, et al. 2015. Genome-wide patterns and properties of *de novo* mutations in humans. Nat Genet 47:822–826.

Frankish A, Diekhans M, Ferreira A-M, Johnson R, Jungreis I, Loveland J, Mudge JM, Sisu C, Wright J, Armstrong J, et al. 2019. GENCODE reference annotation for the human and mouse genomes. Nucleic Acids Res 47:766–773.

Friedlander E, Steinrücken M. 2022. A numerical framework for genetic hitchhiking in populations of variable size. Genetics 220:iyac012.

Galtier N. 2016. Adaptive protein evolution in animals and the effective population size hypothesis. PLoS Genet 12:e1005774.

Gillespie JH. 1984. Molecular evolution over the mutational landscape. Evolution 38:1116.

Gravel S, Henn BM, Gutenkunst RN, Indap AR, Marth GT, Clark AG, Yu F, Gibbs RA, The 1000 Genomes Project, Bustamante CD, et al. 2011. Demographic history and rare allele sharing among human populations. Proc Natl Acad Sci USA 108:11983–11988.

Griffiths RC, Marjoram P. 1997. An ancestral recombination graph. In: Progress in population genetics and human evolution. Vol. 87. New York, NY: Springer New York. p. 257–270.

Gutenkunst RN, Hernandez RD, Williamson SH, Bustamante CD. 2009. Inferring the joint demographic history of multiple populations from multidimensional SNP frequency data. PLoS Genet 5:e1000695.

Haldane JBS. 1919. The combination of linkage values, and the calculation of distances between the loci of linked factors. J Genet 8:299–309.

Haller BC, Messer PW. 2017. asymptoticMK: a web-based tool for the asymptotic McDonald–Kreitman test. G3 7:1569–1575.

Haller BC, Messer PW. 2023. SLiM 4: multispecies eco-evolutionary modeling. Am Nat 201:127–139.

Halligan DL, Kousathanas A, Ness RW, Harr B, Eöry L, Keane TM, Adams DJ, Keightley PD. 2013. Contributions of protein-coding and regulatory change to adaptive molecular evolution in murid rodents. PLoS Genet 9:e1003995.

Harris K, Nielsen R. 2013. Inferring demographic history from a spectrum of shared haplotype lengths. PLoS Genet 9:e1003521.

Hartfield M, Bataillon T. 2020. Selective sweeps under dominance and inbreeding. G3 10:1063–1075.

Hawks J, Hunley K, Lee S-H, Wolpoff M. 2000. Population bottlenecks and Pleistocene human evolution. Mol Biol Evol 17:2–22.

Hayes BJ, Visscher PM, McPartlan HC, Goddard ME. 2003. Novel multilocus measure of linkage disequilibrium to estimate past effective population size. Genome Res 13:635–643.

Hedrick PW. 1980. Hitchhiking: a comparison of linkage and partial selfing. Genetics 94:791–808.

Hoggart CJ, Chadeau-Hyam M, Clark TG, Lampariello R, Whittaker JC, De Iorio M, Balding DJ. 2007. Sequence-level population simulations over large genomic regions. Genetics 177:1725–1731.

Howell AA, Terbot JW, Soni V, Johri P, Jensen JD, Pfeifer SP. 2023. Developing an appropriate evolutionary baseline model for the study of human cytomegalovirus. Genome Biol Evol 15:evad059.

Hu W, Hao Z, Du P, Di Vincenzo F, Manzi G, Cui J, Fu Y-X, Pan Y-H, Li H. 2023. Genomic inference of a severe human bottleneck during the Early to Middle Pleistocene transition. Science 381:979–984.

Hudson RR. 1991. Gene genealogies and the coalescent process. In: Oxford Surveys in Evolutionary Biology. Vol. 7. New York, NY: Oxford University Press. p. 1–44.

Huff CD, Xing J, Rogers AR, Witherspoon D, Jorde LB. 2010. Mobile elements reveal small population size in the ancient ancestors of *Homo sapiens*. Proc Natl Acad Sci USA 107:2147–2152.

Ignatieva A, Favero M, Koskela J, Sant J, Myers SR. 2023. The distribution of branch duration and detection of inversions in ancestral recombination graphs. Available from: http://biorxiv.org/lookup/doi/10.1101/2023.07.11.548567

Ishigohoka J, Liedvogel M. 2024. High-recombining genomic regions affect demography inference. Available from: http://biorxiv.org/lookup/doi/10.1101/2024.02.05.579015

Jensen JD, Payseur BA, Stephan W, Aquadro CF, Lynch M, Charlesworth D, Charlesworth B. 2019. The importance of the Neutral Theory in 1968 and 50 years on: A response to Kern and Hahn 2018. Evolution 73:111–114.

Jensen JD, Thornton KR, Andolfatto P. 2008. An approximate Bayesian estimator suggests strong, recurrent selective sweeps in *Drosophila*. PLoS Genet 4:e1000198.

Jensen-Seaman MI, Furey TS, Payseur BA, Lu Y, Roskin KM, Chen C-F, Thomas MA, Haussler D, Jacob HJ. 2004. Comparative recombination rates in the rat, mouse, and human genomes. Genome Res 14:528–538.

Johri P, Aquadro CF, Beaumont M, Charlesworth B, Excoffier L, Eyre-Walker A, Keightley PD, Lynch M, McVean G, Payseur BA, et al. 2022. Recommendations for improving statistical inference in population genomics. PLoS Biol 20:e3001669.

Johri P, Charlesworth B, Jensen JD. 2020. Toward an evolutionarily appropriate null model: jointly inferring demography and purifying selection. Genetics 215:173–192.

Johri P, Eyre-Walker A, Gutenkunst RN, Lohmueller KE, Jensen JD. 2022. On the prospect of achieving accurate joint estimation of selection with population history. Genome Biol Evol 14:evac088.

Johri P, Pfeifer SP, Jensen JD. 2023. Developing an evolutionary baseline model for humans: jointly inferring purifying selection with population history. Mol Biol Evol 40:msad100.

Johri P, Riall K, Becher H, Excoffier L, Charlesworth B, Jensen JD. 2021. The impact of purifying and background selection on the inference of population history: problems and prospects. Mol Biol Evol 38:2986–3003.

Jouganous J, Long W, Ragsdale AP, Gravel S. 2017. Inferring the joint demographic history of multiple populations: beyond the diffusion approximation. Genetics 206:1549– 1567.

Joyce P, Rokyta DR, Beisel CJ, Orr HA. 2008. A general extreme value theory model for the adaptation of DNA sequences under strong selection and weak mutation. Genetics 180:1627–1643.

Kamm J, Terhorst J, Durbin R, Song YS. 2020. Efficiently inferring the demographic history of many populations with allele count data. J Am Stat Assoc 115:1472–1487.

Kapopoulou A, Pfeifer SP, Jensen JD, Laurent S. 2018. The demographic history of African *Drosophila melanogaster*. Genome Biol Evol 10:2338–2342.

Keightley PD, Campos JL, Booker TR, Charlesworth B. 2016. Inferring the frequency spectrum of derived variants to quantify adaptive molecular evolution in protein-coding genes of *Drosophila melanogaster*. Genetics 203:975–984.

Keightley PD, Ness RW, Halligan DL, Haddrill PR. 2014. Estimation of the spontaneous mutation rate per nucleotide site in a *Drosophila melanogaster* full-sib family. Genetics 196:313–320.

Kelleher J, Wong Y, Wohns AW, Fadil C, Albers PK, McVean G. 2019. Inferring whole-genome histories in large population datasets. Nat Genet 51:1330–1338.

Kim Y. 2006. Allele frequency distribution under recurrent selective sweeps. Genetics 172:1967–1978.

Kim Y, Wiehe T. 2008. Simulation of DNA sequence evolution under models of recent directional selection. Brief Bioinformatics 10:84–96.

Kong A, Frigge ML, Masson G, Besenbacher S, Sulem P, Magnusson G, Gudjonsson SA, Sigurdsson A, Jonasdottir Aslaug, Jonasdottir Adalbjorg, et al. 2012. Rate of *de novo* mutations and the importance of father’s age to disease risk. Nature 488:471–475.

Kong A, Thorleifsson G, Gudbjartsson DF, Masson G, Sigurdsson A, Jonasdottir Aslaug, Walters GB, Jonasdottir Adalbjorg, Gylfason A, Kristinsson KTh, et al. 2010. Fine-scale recombination rate differences between sexes, populations and individuals. Nature 467:1099–1103.

Lange JD, Pool JE. 2018. Impacts of recurrent hitchhiking on divergence and demographic inference in *Drosophila*. Genome Biol Evol 10:1882–1891.

Laurent SJY, Werzner A, Excoffier L, Stephan W. 2011. Approximate Bayesian analysis of *Drosophila melanogaster* polymorphism data reveals a recent colonization of southeast Asia. Mol Biol Evol 28:2041–2051.

Lewanski AL, Grundler MC, Bradburd GS. 2024. The era of the ARG: An introduction to ancestral recombination graphs and their significance in empirical evolutionary genomics. PLoS Genet 20:e1011110.

Li H, Durbin R. 2011. Inference of human population history from individual whole-genome sequences. Nature 475:493–496.

Li N, Stephens M. 2003. Modeling linkage disequilibrium and identifying recombination hotspots using single-nucleotide polymorphism data. Genetics 165:2213–2233.

Lindsay SJ, Rahbari R, Kaplanis J, Keane T, Hurles ME. 2019. Similarities and differences in patterns of germline mutation between mice and humans. Nat Commun 10:4053.

Liu X, Fu Y-X. 2015. Exploring population size changes using SNP frequency spectra. Nat Genet 47:555–559.

Long M, Deutsch M. 1999. Intron—exon structures of eukaryotic model organisms. Nucleic Acids Res 27:3219–3228.

Mackay TFC, Richards S, Stone EA, Barbadilla A, Ayroles JF, Zhu D, Casillas S, Han Y, Magwire MM, Cridland JM, et al. 2012. The *Drosophila melanogaster* genetic reference panel. Nature 482:173–178.

Mao Y, Catacchio CR, Hillier LW, Porubsky D, Li R, Sulovari A, Fernandes JD, Montinaro F, Gordon DS, Storer JM, et al. 2021. A high-quality bonobo genome refines the analysis of hominid evolution. Nature 594:77–81.

Marchi N, Schlichta F, Excoffier L. 2021. Demographic inference. Current Biol 31:R276– R279.

Maynard Smith J, Haigh J. 1974. The hitch-hiking effect of a favourable gene. Genet Res 23:23–35.

Mayr C. 2016. Evolution and biological roles of alternative 3′UTRs. Trends Cell Biol 26:227–237.

Mazet O, Rodríguez W, Grusea S, Boitard S, Chikhi L. 2016. On the importance of being structured: instantaneous coalescence rates and human evolution—lessons for ancestral population size inference. Heredity 116:362–371.

McDonald JH, Kreitman M. 1991. Adaptive protein evolution at the *Adh* locus in *Drosophila*. Nature 351:652–654.

McManus KF, Kelley JL, Song S, Veeramah KR, Woerner AE, Stevison LS, Ryder OA, Ape Genome Project G, Kidd JM, Wall JD, et al. 2015. Inference of gorilla demographic and selective history from whole-genome sequence data. Mol Biol Evol 32:600–612.

McVean GAT, Cardin NJ. 2005. Approximating the coalescent with recombination. Philos Trans R Soc B 360:1387–1393.

Mellars P. 2006. Why did modern human populations disperse from Africa *ca.* 60,000 years ago: a new model. Proc Natl Acad Sci USA 103:9381–9386.

Messer PW, Petrov DA. 2013. Frequent adaptation and the McDonald–Kreitman test. Proc Natl Acad Sci USA 110:8615–8620.

Mignone F, Gissi C, Liuni S, Pesole G. 2002. Untranslated regions of mRNAs. Genome Biol 3:1–10.

Minadakis N, Williams H, Horvath R, Caković D, Stritt C, Thieme M, Bourgeois Y, Roulin AC. 2023. The demographic history of the wild crop relative *Brachypodium distachyon* is shaped by distinct past and present ecological niches. Peer Community J 3:e84.

Moorjani P, Hellenthal G. 2023. Methods for assessing population relationships and history using genomic data. Annu Rev Genomics Hum Genet 24:305–332.

Murphy DA, Elyashiv E, Amster G, Sella G. 2023. Broad-scale variation in human genetic diversity levels is predicted by purifying selection on coding and non-coding elements. eLife 12:e76065.

Nater A, Greminger MP, Arora N, Van Schaik CP, Goossens B, Singleton I, Verschoor EJ, Warren KS, Krützen M. 2015. Reconstructing the demographic history of orangutans using approximate Bayesian computation. Mol Ecol 24:310–327.

Nicolaisen LE, Desai MM. 2013. Distortions in genealogies due to purifying selection and recombination. Genetics 195:221–230.

Nielsen R, Akey JM, Jakobsson M, Pritchard JK, Tishkoff S, Willerslev E. 2017. Tracing the peopling of the world through genomics. Nature 541:302–310.

Nielsen R, Hellmann I, Hubisz M, Bustamante C, Clark AG. 2007. Recent and ongoing selection in the human genome. Nat Rev Genet 8:857–868.

Noble LM, Yuen J, Stevens L, Moya N, Persaud R, Moscatelli M, Jackson JL, Zhang G, Chitrakar R, Baugh LR, et al. 2021. Selfing is the safest sex for *Caenorhabditis tropicalis*. eLife 10:e62587.

Nordborg M. 2000. Linkage disequilibrium, gene trees and selfing: an ancestral recombination graph with partial self-fertilization. Genetics 154:923–929.

Nordborg M, Donnelly P. 1997. The coalescent process with selfing. Genetics 146:1185– 1195.

Novo I, Santiago E, Caballero A. 2022. The estimates of effective population size based on linkage disequilibrium are virtually unaffected by natural selection. PLoS Genet 18:e1009764.

Nurk S, Koren S, Rhie A, Rautiainen M, Bzikadze AV, Mikheenko A, Vollger MR, Altemose N, Uralsky L, Gershman A, et al. 2022. The complete sequence of a human genome. Science 376:44–53.

Orr HA. 2003. The distribution of fitness effects among beneficial mutations. Genetics 163:1519–1526.

Phifer-Rixey M, Bonhomme F, Boursot P, Churchill GA, Pialek J, Tucker PK, Nachman MW. 2012. Adaptive evolution and effective population size in wild house mice. Mol Biol Evol 29:2949–2955.

Piovesan A, Antonaros F, Vitale L, Strippoli P, Pelleri MC, Caracausi M. 2019. Human protein-coding genes and gene feature statistics in 2019. BMC Res Notes 12:315.

Pope NS, Singh A, Childers AK, Kapheim KM, Evans JD, López-Uribe MM. 2023. The expansion of agriculture has shaped the recent evolutionary history of a specialized squash pollinator. Proc Natl Acad Sci USA 120:e2208116120.

Pouyet F, Aeschbacher S, Thiéry A, Excoffier L. 2018. Background selection and biased gene conversion affect more than 95% of the human genome and bias demographic inferences. eLife 7:e36317.

Prado-Martinez J, Sudmant PH, Kidd JM, Li H, Kelley JL, Lorente-Galdos B, Veeramah KR, Woerner AE, O’Connor TD, Santpere G, et al. 2013. Great ape genetic diversity and population history. Nature 499:471–475.

Ragsdale AP, Gravel S. 2019. Models of archaic admixture and recent history from two-locus statistics. PLoS Genet 15:e1008204.

Ragsdale AP, Gutenkunst RN. 2017. Inferring demographic history using two-locus statistics. Genetics 206:1037–1048.

Ragsdale AP, Weaver TD, Atkinson EG, Hoal EG, Möller M, Henn BM, Gravel S. 2023. A weakly structured stem for human origins in Africa. Nature 617:755–763.

Rands CM, Meader S, Ponting CP, Lunter G. 2014. 8.2% of the human genome is constrained: variation in rates of turnover across functional element classes in the human lineage. PLoS Genet 10:e1004525.

Rasmussen MD, Hubisz MJ, Gronau I, Siepel A. 2014. Genome-wide inference of ancestral recombination graphs. PLoS Genet 10:e1004342.

Rhie A, Nurk S, Cechova M, Hoyt SJ, Taylor DJ, Altemose N, Hook PW, Koren S, Rautiainen M, Alexandrov IA, et al. 2023. The complete sequence of a human Y chromosome. Nature 621:344–354.

Santiago E, Novo I, Pardiñas AF, Saura M, Wang J, Caballero A. 2020. Recent demographic history inferred by high-resolution analysis of linkage disequilibrium. Mol Biol Evol 37:3642–3653.

dos Santos G, Schroeder AJ, Goodman JL, Strelets VB, Crosby MA, Thurmond J, Emmert DB, Gelbart WM, the FlyBase Consortium. 2015. FlyBase: introduction of the *Drosophila melanogaster* Release 6 reference genome assembly and large-scale migration of genome annotations. Nucleic Acids Res 43:690–697.

Sayers EW, Bolton EE, Brister JR, Canese K, Chan J, Comeau DC, Farrell CM, Feldgarden M, Fine AM, Funk K, et al. 2023. Database resources of the National Center for Biotechnology Information in 2023. Nucleic Acids Res 51:D29–D38.

Schiffels S, Durbin R. 2014. Inferring human population size and separation history from multiple genome sequences. Nat Genet 46:919–925.

Schiffels S, Wang K. 2020. MSMC and MSMC2: the multiple sequentially Markovian coalescent. In: Statistical Population Genomics. Vol. 2090. Methods in Molecular Biology. New York, NY: Springer US. p. 147–166.

Schlebusch CM, Malmström H, Günther T, Sjödin P, Coutinho A, Edlund H, Munters AR, Vicente M, Steyn M, Soodyall H, et al. 2017. Southern African ancient genomes estimate modern human divergence to 350,000 to 260,000 years ago. Science 358:652–655.

Schmidt JM, De Manuel M, Marques-Bonet T, Castellano S, Andrés AM. 2019. The impact of genetic adaptation on chimpanzee subspecies differentiation. PLoS Genet 15:e1008485.

Schrider DR, Shanku AG, Kern AD. 2016. Effects of linked selective sweeps on demographic inference and model selection. Genetics 204:1207–1223.

Seetharaman S, Jain K. 2014. Adaptive walks and distribution of beneficial fitness effects. Evolution 68:965–975.

Sellinger TPP, Abu-Awad D, Tellier A. 2021. Limits and convergence properties of the sequentially Markovian coalescent. Mol Ecol Res 21:2231–2248.

Sheehan S, Song YS. 2016. Deep learning for population genetic inference. PLoS Comput Biol 12:e1004845.

Sherry ST, Harpending HC, Batzer MA, Stoneking M. 1997. *Alu* Evolution in human populations: using the coalescent to estimate effective population size. Genetics 147:1977–1982.

Siepel A, Bejerano G, Pedersen JS, Hinrichs AS, Hou M, Rosenbloom K, Clawson H, Spieth J, Hillier LW, Richards S, et al. 2005. Evolutionarily conserved elements in vertebrate, insect, worm, and yeast genomes. Genome Res 15:1034–1050.

Smith ML, Hahn MW. 2023. Selection leads to false inferences of introgression using popular methods. Available from: http://biorxiv.org/lookup/doi/10.1101/2023.10.27.564394

Soni V, Johri P, Jensen JD. 2023. Evaluating power to detect recurrent selective sweeps under increasingly realistic evolutionary null models. Evolution 77:2113–2127.

Soni V, Pfeifer SP, Jensen JD. 2024. The effects of mutation and recombination rate heterogeneity on the inference of demography and the distribution of fitness effects. Genome Biol Evol 16:evae004.

Speidel L, Forest M, Shi S, Myers SR. 2019. A method for genome-wide genealogy estimation for thousands of samples. Nat Genet 51:1321–1329.

Strütt S, Sellinger T, Glémin S, Tellier A, Laurent S. 2023. Joint inference of evolutionary transitions to self-fertilization and demographic history using whole-genome sequences. eLife 12:e82384.

Tataru P, Bataillon T. 2019. polyDFEv2.0: testing for invariance of the distribution of fitness effects within and across species. Bioinformatics 35:2868–2869.

Terbot JW, Johri P, Liphardt SW, Soni V, Pfeifer SP, Cooper BS, Good JM, Jensen JD. 2023. Developing an appropriate evolutionary baseline model for the study of SARS-CoV-2 patient samples. Hobman TC, editor. PLoS Pathog 19:e1011265.

Terhorst J, Kamm JA, Song YS. 2017. Robust and scalable inference of population history from hundreds of unphased whole genomes. Nat Genet 49:303–309.

Teshima KM, Coop G, Przeworski M. 2006. How reliable are empirical genomic scans for selective sweeps? Genome Res 16:702–712.

Teshima KM, Przeworski M. 2006. Directional positive selection on an allele of arbitrary dominance. Genetics 172:713–718.

Teterina AA, Willis JH, Lukac M, Jovelin R, Cutter AD, Phillips PC. 2023. Genomic diversity landscapes in outcrossing and selfing *Caenorhabditis nematodes*. PLoS Genet 19:e1010879.

The International HapMap Consortium. 2005. A haplotype map of the human genome. Nature 437:1299–1320.

Thornton K, Andolfatto P. 2006. Approximate Bayesian inference reveals evidence for a recent, severe bottleneck in a Netherlands population of *Drosophila melanogaster*. Genetics 172:1607–1619.

Thornton KR, Jensen JD. 2007. Controlling the false-positive rate in multilocus genome scans for selection. Genetics 175:737–750.

Upadhya G, Steinrücken M. 2022. Robust inference of population size histories from genomic sequencing data. PLoS Comput Biol 18:e1010419.

Uricchio LH, Hernandez RD. 2014. Robust forward simulations of recurrent hitchhiking. Genetics 197:221–236.

Uricchio LH, Petrov DA, Enard D. 2019. Exploiting selection at linked sites to infer the rate and strength of adaptation. Nat Ecol Evol 3:977–984.

Wohns AW, Wong Y, Jeffery B, Akbari A, Mallick S, Pinhasi R, Patterson N, Reich D, Kelleher J, McVean G. 2022. A unified genealogy of modern and ancient genomes. Science 375:eabi8264.

Wong Y, Ignatieva A, Koskela J, Gorjanc G, Wohns AW, Kelleher J. 2023. A general and efficient representation of ancestral recombination graphs. Available from: http://biorxiv.org/lookup/doi/10.1101/2023.11.03.565466

Wright SI, Kalisz S, Slotte T. 2013. Evolutionary consequences of self-fertilization in plants. Philos Trans R Soc B 280:20130133.

Zeng K, Charlesworth B. 2011. The joint effects of background selection and genetic recombination on local gene genealogies. Genetics 189:251–266.

Zhang BC, Biddanda A, Gunnarsson ÁF, Cooper F, Palamara PF. 2023. Biobank-scale inference of ancestral recombination graphs enables genealogical analysis of complex traits. Nat Genet 55:768–776.

Zhen Y, Huber CD, Davies RW, Lohmueller KE. 2021. Greater strength of selection and higher proportion of beneficial amino acid changing mutations in humans compared with mice and *Drosophila melanogaster*. Genome Res 31:110–120.

