## Supplemental file for "Biases in ARG-based inference of historical population size in populations experiencing selection" (from another version of this preprint: version 1, 2024-04-26)

### Supplementary Figures, Notes, and Tables

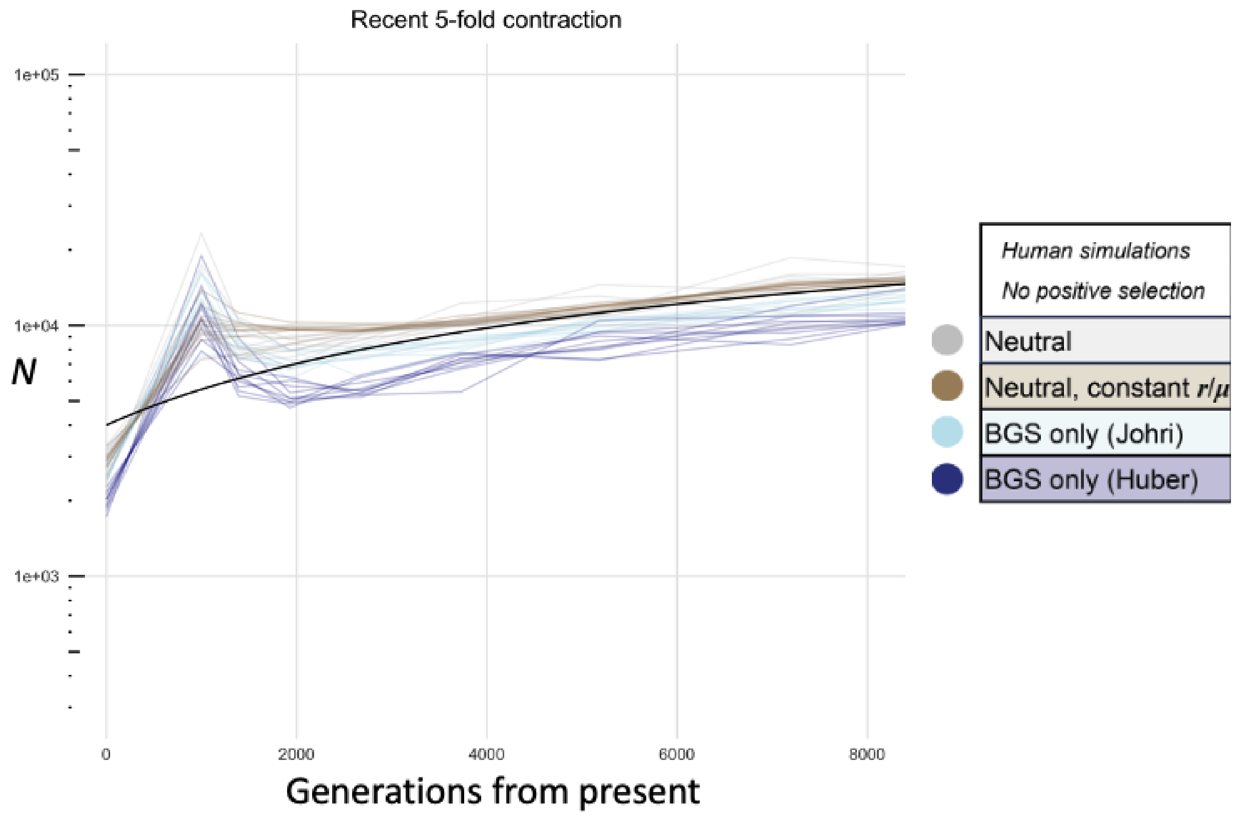

**Figure S1.** Demography inferred by *Relate* for human-like simulated data in most recent 8000 generations prior to present, under the recent 5-fold population contraction scenario (identical to Figure 1C with truncated x-axis). Note the spike in population size (*N*) estimated approximately 1000 generations ago for all conditions.

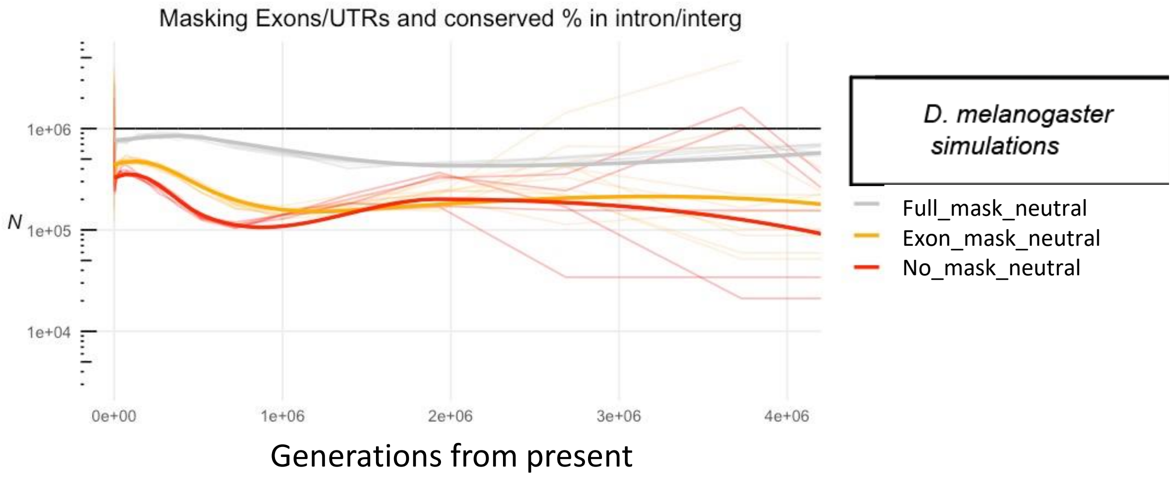

**Figure S2.** Demography inferred by *Relate* for *D. melanogaster*-like simulated data under constant population size with no selection present (same as Figure 4A). All three conditions use the same input VCF, which had variants in "conserved" regions (exons, UTRs, 43% of introns, 35% of intergenic) removed totalling 48.7% of all sites, though with differing genomic masks provided. No genomic mask provided (red), masking variant sites only in regions experiencing selection (orange; 48.7% of variant sites), masking variant sites experiencing selection as well as all sites (variant and nonvariant) in "conserved" regions (grey; 48.7% of variant and nonvariant sites).

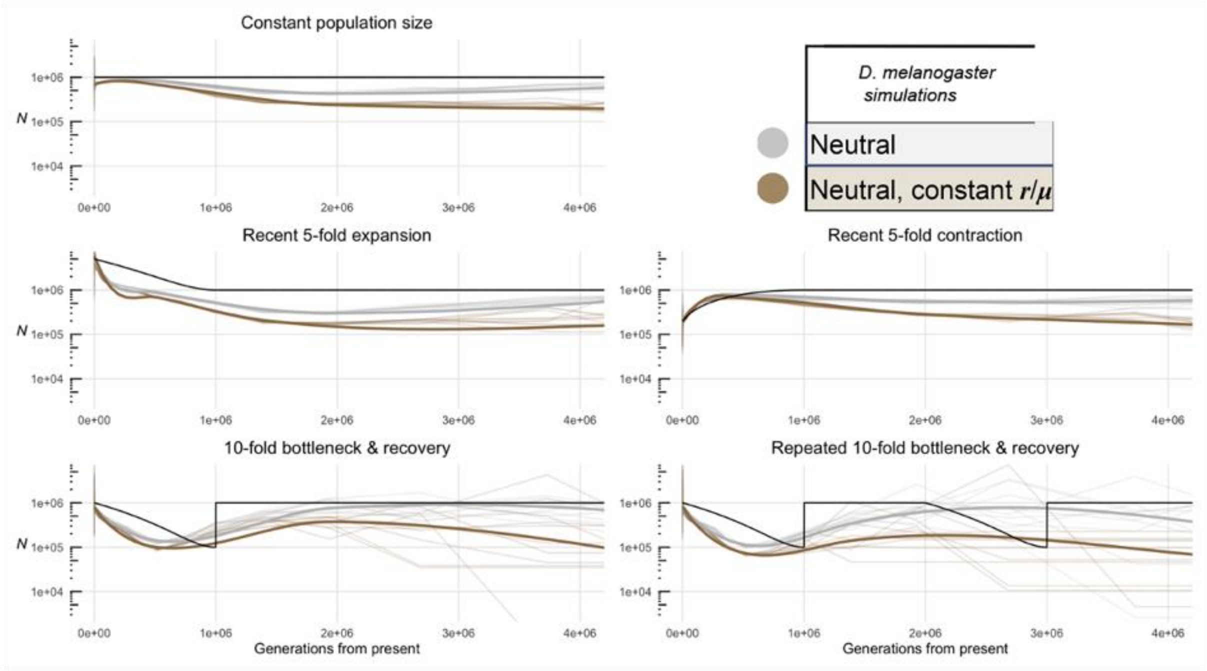

**Figure S3.** Demography inferred by *Relate* for *D. melanogaster*-like simulated data under constant population size with no selection present (same as Figure 4A) with variable recombination and mutation rates simulated across the genome (grey) and with constant mutation and recombination rates simulated across the genome (gold). In both cases, accurate recombination rate maps were provided to *Relate*.

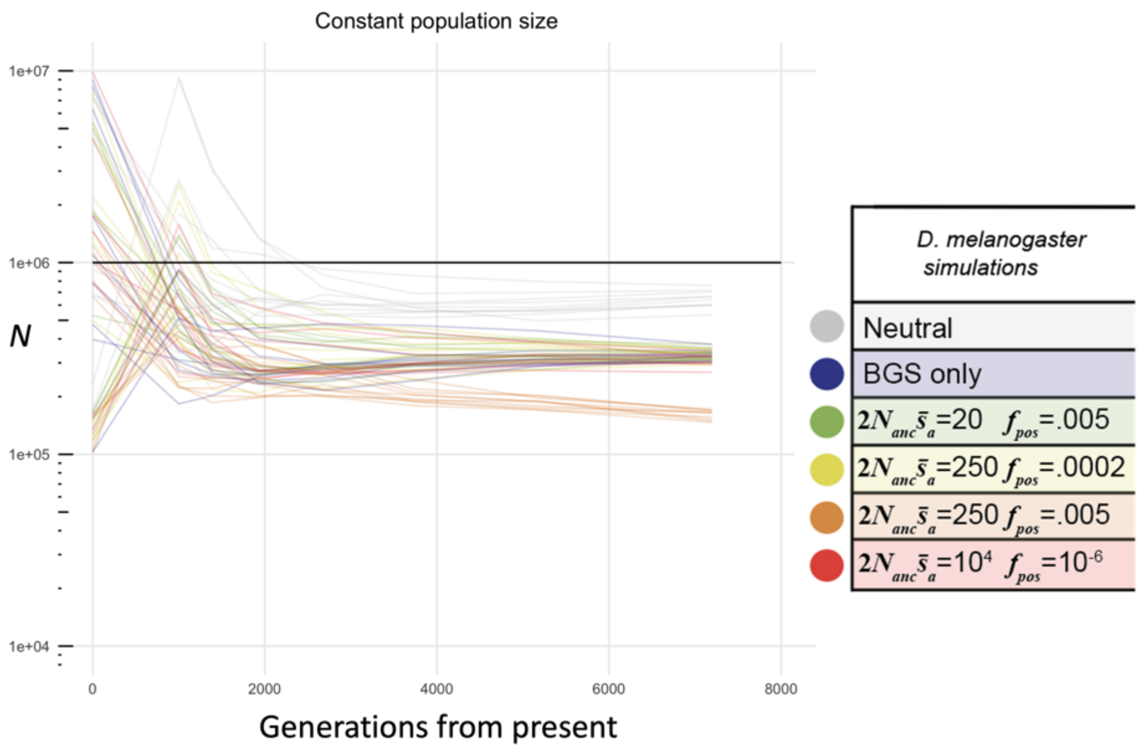

**Figure S4.** Demography inferred by *Relate* for *Drosophila*-like simulated data in most recent 8000 generations prior to present, under constant population size (identical to Figure 4A with truncated X-axis). Note the extreme stochasticity in all conditions in the most recent ~2500 generations.

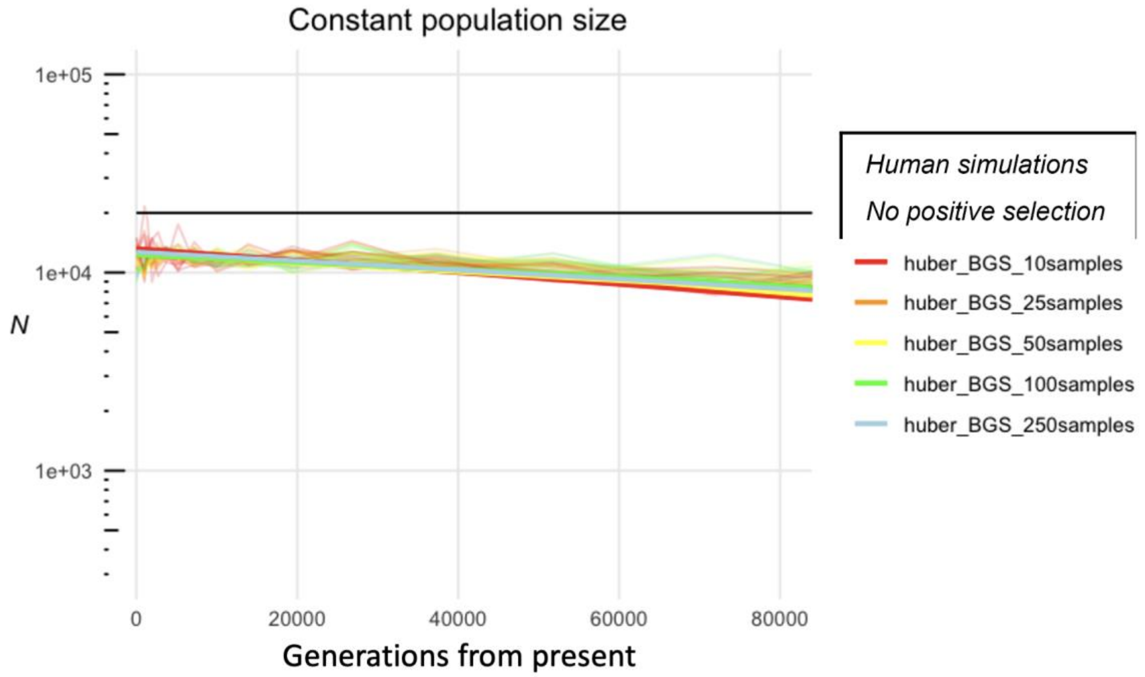

**Figure S5.** Demography inferred by *Relate* for human-like simulated data under constant population size using different number of samples (10 to 250) of diploid individuals as input to *Relate*. Simulations had variable recombination and mutation rates and experienced BGS (DFE from Huber et al. 2017), with no positive selection (same as BGS only in Figure 1A).

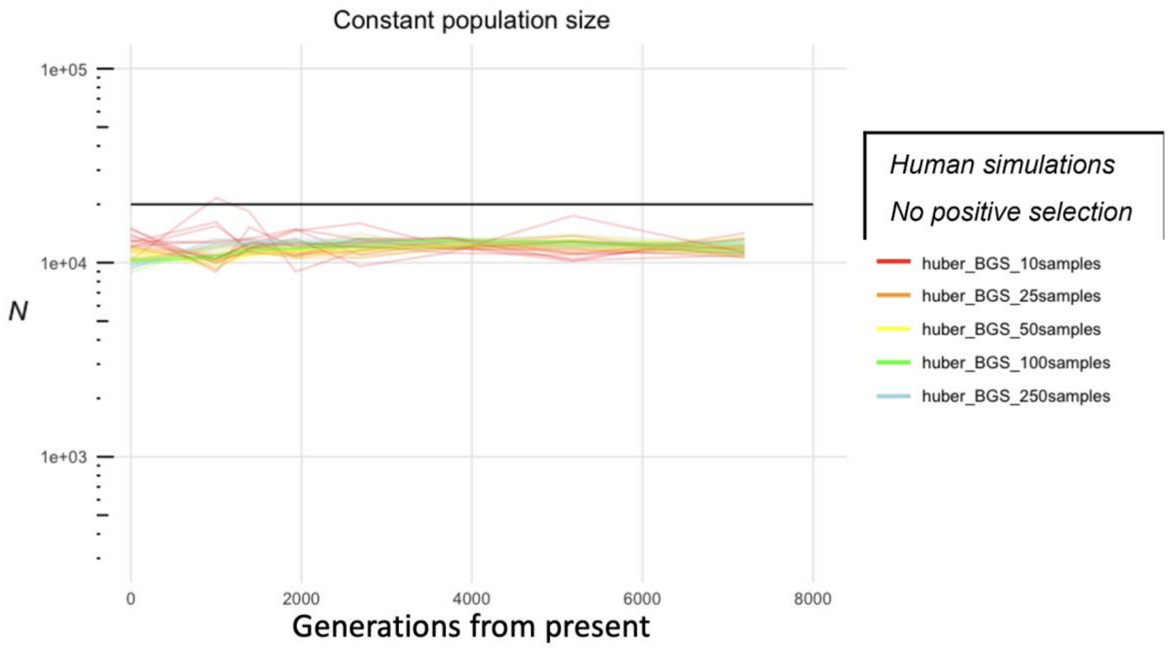

**Figure S6.** Demography inferred by *Relate* for human-like simulated data under constant population size in the most recent 8000 generations using different number of samples (10 to 250) of diploid individuals as input to *Relate*. Simulations had variable recombination and mutation rates and experienced BGS (DFE from Huber et al. 2017), with no positive selection (same as BGS only in Figure 1A).

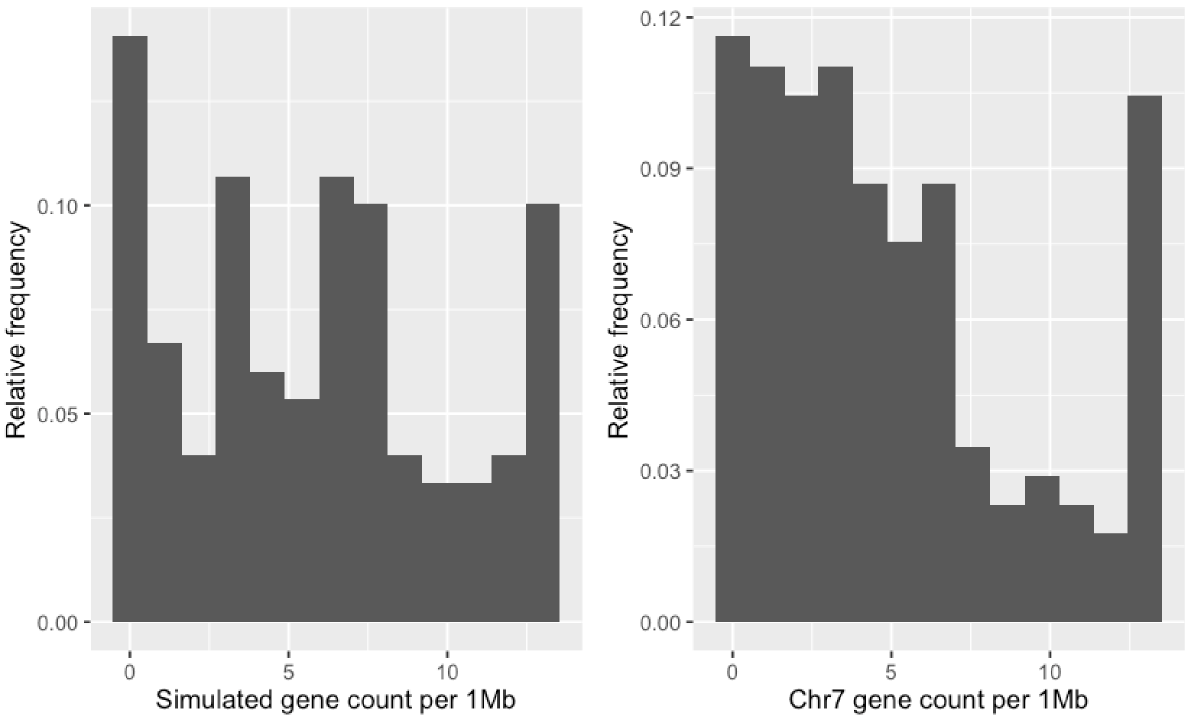

**Figure S7.** Distribution of different gene counts in each 1Mb bin across the simulated 138.6 Mb human chromosome (left) and for human chromosome 7 (right). The counts for human chromosome 7 (right) are truncated at 13, which is the maximum number of genes of the simulated length that fit in a 1Mb region without overlap.

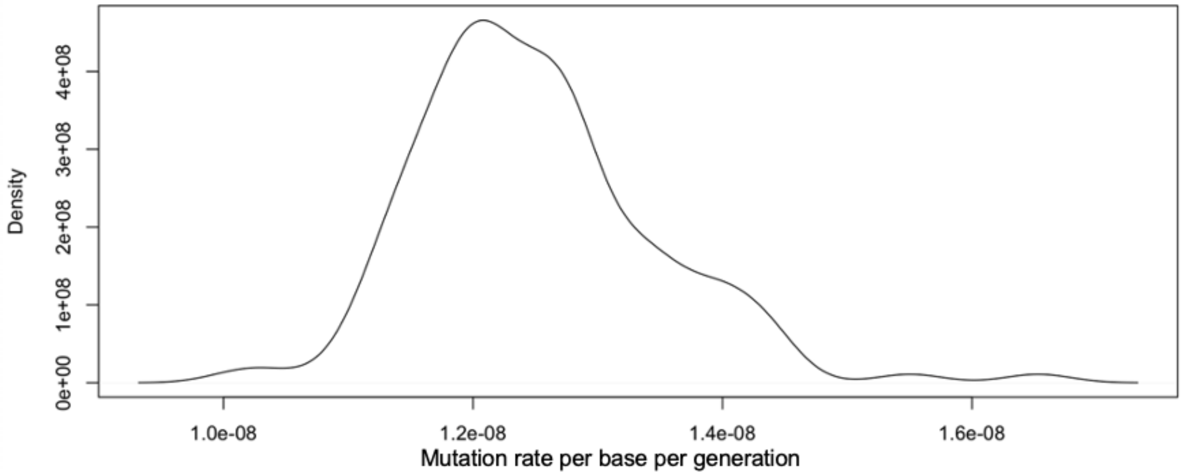

**Figure S8.** Probability density distribution of mutation rates in the mutation rate map from human chromosome 7 (Francioli et al. 2015) used for human simulations, after scaling all values by 1.266 (given the mean mutation rate on chromosome 7 is 0.99e-8 per site per generation) to reach a mean per base per generation mutation rate of 1.25e-8.

**Supplementary Note S1.** Impacts of not supplying an effective genomic mask file to *Relate* for demographic inference with a proportional reduction in genomic distance matching the fraction of variant sites removed from analysis.

When removing sites from the variant file (VCF) provided to *Relate*, such as those under selection in our case or *e.g.* empirical data with low mapping quality, we find that it is essential to provide an accurate mask file to *Relate* that not only indicates which individual sites are masked, but often more critically, masks a fraction of genetic length of the region proportional to the fraction of variants removed from the analysis.

This can be illustrated by observing what happens when, under strict neutrality, half the variant sites in the VCF are removed from the analysis (as was done for *Drosophila* due to ~48% of sites experiencing selection) and how demographic inference results change when providing different genomic masks (Figure S2). Notice that unless a similar proportion of sites removed from the VCF is reflected in the genetic distance masked by the input mask file, historical *N* estimates are lowered, high stochasticity is present between replicates, and recent population expansion is apparent, all of which is similar to the linked effects of selection we report in the main text (Figure 4).

As a result, in our simulations we built genomic masks not only for variant sites under direct selection, but also for the proportion of sites experiencing selection (ensuring neutral variant sites were not masked).

**Supplementary Note S2.** Reductions of diversity at neutral sites compared to all sites for *D. melanogaster* and *E. pruinosa* populations.

The squash bee (*Eucera pruinosa*) genome has a less compact genome than *D. melanogaster* (*e.g.,* ~14000 genes in ~180 Mb for *D. melanogaster* compared to ~8000 genes in ~400 Mb *E. pruinosa*; dos Santos et al. 2015; Pope et al. 2023), however, the relative reductions of diversity at neutral sites due to linked effects of selection in the MO, CO, and PA squash bee populations (Pope et al. 2023) matched the reductions of diversity at neutral sites compared to all sites in our simulations of *D. melanogaster* populations experiencing background selection only. This could suggest similar linked effects of selection in neutral regions for some *E. pruinosa* populations to our BGS simulations (Figure 4) which may have caused mis-inference, assuming *coaldecoder* is similarly sensitive to BGS as *Relate*:

"Observed and expected summary statistics from five populations of *Eucera pruinosa*. Observed statistics are calculated using the entire accessible genome ("all") or a high-diversity subset of intervals assumed to be free of selective sweeps ("neutral", see main text)." (Caption and results from Table S2; Pope et al. 2023):

| *Population* | $\pi_{all}/\pi_{neutral}$ |
|---|---|
| AZ | 0.89 |
| CO | 0.77 |
| MX | 0.94 |
| PA | 0.63 |
| MO | 0.66 |

The $\pi_{all}/\pi_{neu}$ for *D. melanogaster* simulations (our study) with background selection only, where $\pi_{neutral}$ was calculated using neutral sites not under selection ($\pi_{neu}$) was the following (*n.b.* Especially relevant are the simulated population expansion and bottleneck scenarios, which lead to similar findings when inferred by *Relate* with BGS present; Figure 4):

| *Demography* | $\pi_{all}/\pi_{neutral}$ |
|---|---|
| Constant | 0.72 |
| Expansion | 0.66 |
| Contraction | 0.69 |
| Bottleneck | 0.75 |
| Repbneck | 0.75 |

**Table S1.** Nucleotide diversity in neutral intergenic/intronic regions ($\pi_{neu}$) for human simulations. "With maps" refers to neutral simulations with realistic heterogeneity of mutation and recombination rates modelled, while "no maps" refers to neutral simulations where mutation and recombination rates were constant. The deleterious DFE that we used to simulate background selection (BGS) is indicated by the study that estimated the DFE in the first column. Parameters of recurrent positive selection employed in our simulations are provided by the proportion of new mutations that are beneficial ($f_{\text{pos}}$) and the mean population-scaled strength of selection ($2N_{anc}\bar{s_a}$).

| BGS | $f_{\text{pos}}$ | $2N_{anc}\bar{s_a}$ | Constant | Expansion | Contraction | Bottleneck | Repbneck |
|---|---|---|---|---|---|---|---|
| None (no maps) | 0 | N/A | 0.00100 | 0.00121 | 0.00075 | 0.00043 | 0.00038 |
| None (with maps) | 0 | N/A | 0.00100 | 0.00121 | 0.00075 | 0.00042 | 0.00038 |
| Johri | 0 | N/A | 0.00085 | 0.00105 | 0.00063 | 0.00037 | 0.00034 |
| Huber | 0 | N/A | 0.00068 | 0.00088 | 0.00047 | 0.00029 | 0.00028 |
| Huber | 0.001 | 10 | 0.00068 | 0.00088 | 0.00047 | 0.00029 | 0.00028 |
| Huber | 0.001 | 100 | 0.00057 | 0.00069 | 0.00040 | 0.00028 | 0.00027 |
| Huber | 0.001 | 500 | 0.00024 | 0.00021 | 0.00020 | 0.00019 | 0.00019 |
| Huber | 0.001 | 1000 | 0.00012 | 0.00009 | 0.00012 | 0.00012 | 0.00012 |
| Huber | 0.01 | 10 | 0.00066 | 0.00084 | 0.00045 | 0.00029 | 0.00028 |
| Huber | 0.01 | 100 | 0.00029 | 0.00030 | 0.00023 | 0.00021 | 0.00021 |
| Huber | 0.001 | 500 | 0.00006 | 0.00005 | 0.00006 | 0.00007 | 0.00007 |
| Huber | 0.01 | 1000 | 0.00003 | 0.00003 | 0.00003 | 0.00003 | 0.00003 |

**Table S2.** The ratio of diversity in neutral regions ($\pi_{neu}$) compared for human simulations with listed parameter combinations, compared to under strict neutrality. "With maps" refers to neutral simulations with realistic heterogeneity of mutation and recombination rates modelled, while "no maps" refers to neutral simulations where mutation and recombination rates were constant. The deleterious DFE that we used to simulate background selection (BGS) is indicated by the study that estimated the DFE in the first column. Parameters of recurrent positive selection employed in our simulations are provided by the proportion of new mutations that are beneficial ($f_{pos}$) and the mean population-scaled strength of selection ($2N_{anc}\bar{s_a}$).

| BGS | $f_{pos}$ | $2N_{anc}\bar{s_a}$ | Constant | Expansion | Contraction | Bottleneck | Repbneck |
|---|---|---|---|---|---|---|---|
| None (no maps) | 0 | N/A | 1.00 | 1.00 | 1.00 | 1.00 | 1.00 |
| None (with maps) | 0 | N/A | 1.00 | 1.00 | 1.00 | 1.00 | 1.00 |
| Johri | 0 | N/A | 0.86 | 0.87 | 0.84 | 0.87 | 0.90 |
| Huber | 0 | N/A | 0.69 | 0.73 | 0.62 | 0.70 | 0.75 |
| Huber | 0.001 | 10 | 0.68 | 0.73 | 0.62 | 0.69 | 0.75 |
| Huber | 0.001 | 100 | 0.58 | 0.57 | 0.53 | 0.65 | 0.71 |
| Huber | 0.001 | 500 | 0.24 | 0.17 | 0.26 | 0.44 | 0.49 |
| Huber | 0.001 | 1000 | 0.12 | 0.07 | 0.16 | 0.29 | 0.32 |
| Huber | 0.01 | 10 | 0.66 | 0.70 | 0.60 | 0.69 | 0.74 |
| Huber | 0.01 | 100 | 0.29 | 0.25 | 0.30 | 0.50 | 0.56 |
| Huber | 0.001 | 500 | 0.07 | 0.04 | 0.08 | 0.16 | 0.17 |
| Huber | 0.01 | 1000 | 0.03 | 0.02 | 0.05 | 0.07 | 0.08 |

**Table S3.** Divergence values in neutral intergenic/intronic regions for human simulations with listed parameter combinations. "With maps" refers to neutral simulations with realistic heterogeneity of mutation and recombination rates modelled, while "no maps" refers to neutral simulations where mutation and recombination rates were constant. The deleterious DFE that we used to simulate background selection (BGS) is indicated by the study that estimated the DFE in the first column. Parameters of recurrent positive selection employed in our simulations are provided by the proportion of new mutations that are beneficial ($f_{pos}$) and the mean population-scaled strength of selection ($2N_{anc}\bar{s_a}$).

| BGS | $f_{pos}$ | $2N_{anc}\bar{s_a}$ | Constant | Expansion | Contraction | Bottleneck | Repbneck |
|---|---|---|---|---|---|---|---|
| None (no maps) | N/A | N/A | 0.00099 | 0.00113 | 0.00140 | 0.00184 | 0.00240 |
| None (with maps) | N/A | N/A | 0.00099 | 0.00113 | 0.00139 | 0.00183 | 0.00239 |
| Johri | 0 | N/A | 0.00099 | 0.00114 | 0.00138 | 0.00175 | 0.00229 |
| Huber | 0 | N/A | 0.00099 | 0.00113 | 0.00139 | 0.00167 | 0.00218 |
| Huber | 0.001 | 10 | 0.00099 | 0.00113 | 0.00138 | 0.00167 | 0.00218 |
| Huber | 0.001 | 100 | 0.00099 | 0.00116 | 0.00135 | 0.00156 | 0.00207 |
| Huber | 0.001 | 500 | 0.00100 | 0.00126 | 0.00127 | 0.00131 | 0.00181 |
| Huber | 0.001 | 1000 | 0.00100 | 0.00127 | 0.00125 | 0.00125 | 0.00175 |
| Huber | 0.01 | 10 | 0.00099 | 0.00114 | 0.00138 | 0.00164 | 0.00215 |
| Huber | 0.01 | 100 | 0.00100 | 0.00122 | 0.00129 | 0.00134 | 0.00184 |
| Huber | 0.001 | 500 | 0.00099 | 0.00125 | 0.00124 | 0.00124 | 0.00174 |
| Huber | 0.01 | 1000 | 0.00099 | 0.00125 | 0.00124 | 0.00124 | 0.00174 |

**Table S4.** Divergence values in coding regions for human simulations with listed parameter combinations. "With maps" refers to neutral simulations with realistic heterogeneity of mutation and recombination rates modelled, while "no maps" refers to neutral simulations where mutation and recombination rates were constant. The deleterious DFE that we used to simulate background selection (BGS) is indicated by the study that estimated the DFE in the first column. Parameters of recurrent positive selection employed in our simulations are provided by the proportion of new mutations that are beneficial ($f_{pos}$) and the mean population-scaled strength of selection ($2N_{anc}\bar{s_a}$).

| BGS | $f_{pos}$ | $2N_{anc}\bar{s_a}$ | Constant | Expansion | Contraction | Bottleneck | Repbneck |
|---|---|---|---|---|---|---|---|
| None (no maps) | 0 | N/A | 0.00100 | 0.00114 | 0.00140 | 0.00185 | 0.00241 |
| None (with maps) | 0 | N/A | 0.00099 | 0.00112 | 0.00138 | 0.00182 | 0.00237 |
| Johri | 0 | N/A | 0.00052 | 0.00060 | 0.00072 | 0.00093 | 0.00124 |
| Huber | 0 | N/A | 0.00052 | 0.00060 | 0.00073 | 0.00089 | 0.00118 |
| Huber | 0.001 | 10 | 0.00054 | 0.00061 | 0.00076 | 0.00092 | 0.00120 |
| Huber | 0.001 | 100 | 0.00060 | 0.00072 | 0.00081 | 0.00092 | 0.00123 |
| Huber | 0.001 | 500 | 0.00085 | 0.00112 | 0.00106 | 0.00106 | 0.00145 |
| Huber | 0.001 | 1000 | 0.00101 | 0.00132 | 0.00124 | 0.00121 | 0.00166 |
| Huber | 0.01 | 10 | 0.00060 | 0.00069 | 0.00083 | 0.00097 | 0.00129 |
| Huber | 0.01 | 100 | 0.00109 | 0.00139 | 0.00137 | 0.00134 | 0.00182 |
| Huber | 0.001 | 500 | 0.00169 | 0.00219 | 0.00208 | 0.00200 | 0.00274 |
| Huber | 0.01 | 1000 | 0.00186 | 0.00240 | 0.00229 | 0.00222 | 0.00303 |

**Table S5.** Ratio of divergence values in coding/neutral regions for human simulations with listed parameter combinations. "With maps" refers to neutral simulations with realistic heterogeneity of mutation and recombination rates modelled, while "no maps" refers to neutral simulations where mutation and recombination rates were constant. The deleterious DFE that we used to simulate background selection (BGS) is indicated by the study that estimated the DFE in the first column. Parameters of recurrent positive selection employed in our simulations are provided by the proportion of new mutations that are beneficial ($f_{\text{pos}}$) and the mean population-scaled strength of selection ($2N_{anc}\bar{s_a}$).

| BGS | $f_{\text{pos}}$ | $2N_{anc}\bar{s_a}$ | Constant | Expansion | Contraction | Bottleneck | Repbneck |
|---|---|---|---|---|---|---|---|
| None (no maps) | 0 | N/A | 1.01 | 1.01 | 1.00 | 1.01 | 1.00 |
| None (with maps) | 0 | N/A | 1.00 | 0.99 | 0.99 | 1.00 | 0.99 |
| Johri | 0 | N/A | 0.52 | 0.53 | 0.52 | 0.53 | 0.54 |
| Huber | 0 | N/A | 0.53 | 0.53 | 0.53 | 0.53 | 0.54 |
| Huber | 0.001 | 10 | 0.54 | 0.54 | 0.55 | 0.55 | 0.55 |
| Huber | 0.001 | 100 | 0.61 | 0.63 | 0.60 | 0.59 | 0.59 |
| Huber | 0.001 | 500 | 0.85 | 0.89 | 0.83 | 0.81 | 0.80 |
| Huber | 0.001 | 1000 | 1.01 | 1.04 | 0.99 | 0.97 | 0.95 |
| Huber | 0.01 | 10 | 0.60 | 0.61 | 0.60 | 0.59 | 0.60 |
| Huber | 0.01 | 100 | 1.09 | 1.13 | 1.06 | 1.00 | 0.99 |
| Huber | 0.01 | 500 | 1.70 | 1.75 | 1.67 | 1.61 | 1.58 |
| Huber | 0.01 | 1000 | 1.87 | 1.92 | 1.85 | 1.78 | 1.74 |

**Table S6.** Alpha values (proportion of substitutions in coding regions that experienced positive selection, scaled to represent nonsynonymous sites) for human simulations with listed parameter combinations. The deleterious DFE that we used to simulate background selection (BGS) is indicated by the study that estimated the DFE in the first column. Parameters of recurrent positive selection employed in our simulations are provided by the proportion of new mutations that are beneficial ($f_{pos}$) and the mean population-scaled strength of selection ($2N_{anc}\bar{s_a}$).

| BGS | $f_{pos}$ | $2N_{anc}\bar{s_a}$ | Constant | Expansion | Contraction | Bottleneck | Repbneck |
|---|---|---|---|---|---|---|---|
| Huber | 0.001 | 10 | 0.033 | 0.033 | 0.030 | 0.023 | 0.021 |
| Huber | 0.001 | 100 | 0.221 | 0.243 | 0.203 | 0.165 | 0.156 |
| Huber | 0.001 | 500 | 0.451 | 0.474 | 0.440 | 0.413 | 0.399 |
| Huber | 0.001 | 1000 | 0.516 | 0.533 | 0.509 | 0.490 | 0.478 |
| Huber | 0.01 | 10 | 0.225 | 0.230 | 0.214 | 0.176 | 0.164 |
| Huber | 0.01 | 100 | 0.633 | 0.648 | 0.624 | 0.594 | 0.580 |
| Huber | 0.01 | 500 | 0.730 | 0.740 | 0.726 | 0.714 | 0.707 |
| Huber | 0.01 | 1000 | 0.736 | 0.744 | 0.732 | 0.723 | 0.717 |

**Table S7.** Summary of parameters used for *D. melanogaster* simulations prior to rescaling, with corresponding published estimates and any methods used to calculate values (see main text for full reference).

| | Pre-scaled values used in simulations | Published | Source |
|---|---|---|---|
| $N_e$ | 1.00E+06 | 1.40E+06 | Keightley et al. 2014 |
| Mean Mutation rate | 3E-9* | 2.80E-09 | Keightley et al. 2014 |
| Mutation rate Std dev/Mean ratio | 0.479^ | 0.479^ | Taken directly from 4-fold divergence (k) with *D. simulans* sd/mean ratio for 2R middle section (Supplementary Table 11 in Mackay et al. 2012) |
| Mean Recombination rate | 1E-8* | 1.33E-8^† | Comeron et al. 2012 |
| CDS length per exon | 314bp | 360bp^ | Calculated from BDGP6 Chr2R GFF3 (dos Santos et al. 2014) |
| 5'UTR length | 266bp | 266bp^ | Calculated from BDGP6 Chr2R GFF3 (dos Santos et al. 2014) |
| 3'UTR length | 382bp | 382bp^ | Calculated from BDGP6 Chr2R GFF3 (dos Santos et al. 2014) |
| Intron length | 999bp | 1000bp^ | Calculated from BDGP6 Chr2R GFF3 (dos Santos et al. 2014) |
| Mean Intergenic length | 2654bp | 2654bp^ | Calculated from BDGP6 Chr2R GFF3 (dos Santos et al. 2014) |
| Exon number | 5 | 5.24^ | Calculated assuming 1570bp, and exon number = intron number + 1 (Deutsch & Long 1999) |

| | | | |
|---|---|---|---|
| Total CDS length per gene | 1570bp | 1568bp^ | Calculated from BDGP6 Chr2R GFF3 (dos Santos et al. 2014) |
| Intron number | 4 | 4.24^ | Calculated assuming 1570bp (Deutsch & Long 1999) |
| Total single gene length | 6214bp | 6215bp^ | Calculated from BDGP6 Chr2R GFF3 (dos Santos et al. 2014) |
| Genes (per 5Mb) | 565 | 565^ | Calculated from BDGP6 Chr2R GFF3 (dos Santos et al. 2014) |
| CDS % of region | 17.70% | 17.68%^ | Calculated from BDGP6 Chr2R GFF3 (dos Santos et al. 2014) |
| Intron Constrained % | 43 | 43 | Siepel et al. 2005 |
| Intergenic Constrained % | 35 | 35 | Siepel et al. 2005 |
| Total Constrained % | 48.7 | 45 (39-53) | Siepel et al. 2005 |
| *$N_e$ was scaled down by a factor of 300, while mutation rate and recombination rate were multiplied by 300 for simulations | | | |
| ^Published values restricted to results for Chromosome 2R | | | |
| †Sex-averaged recombination rate was obtained by halving the 2R rate provided by Comeron et al. (2012) (2.66E-8) | | | |

**Table S8.** Nucleotide diversity in neutral intergenic/intronic regions ($\pi_{neu}$) for *D. melanogaster* simulations. "With maps" refers to neutral simulations with realistic heterogeneity of mutation and recombination rates modelled, while "no maps" refers to neutral simulations where mutation and recombination rates were constant. The deleterious DFE that we used to simulate background selection (BGS) is indicated by the study that estimated the DFE in the first column. Parameters of recurrent positive selection employed in our simulations are provided by the proportion of new mutations that are beneficial ($f_{pos}$) and the mean population-scaled strength of selection ($2N_{anc}\bar{s_a}$).

| BGS | $f_{pos}$ | $2N_{anc}\bar{s_a}$ | Constant | Expansion | Contraction | Bottleneck | Repbneck |
|---|---|---|---|---|---|---|---|
| None (no maps) | N/A | N/A | 0.01195 | 0.01451 | 0.00904 | 0.00512 | 0.00460 |
| None (with maps) | N/A | N/A | 0.01172 | 0.01423 | 0.00885 | 0.00500 | 0.00448 |
| Johri | 0 | N/A | 0.00472 | 0.00689 | 0.00270 | 0.00241 | 0.00241 |
| Johri | 0.0002 | 20 | 0.00473 | 0.00688 | 0.00270 | 0.00239 | 0.00240 |
| Johri | 0.0002 | 250 | 0.00405 | 0.00530 | 0.00242 | 0.00229 | 0.00229 |
| Johri | 0.005 | 20 | 0.00456 | 0.00655 | 0.00262 | 0.00239 | 0.00239 |
| Johri | 0.005 | 250 | 0.00138 | 0.00129 | 0.00106 | 0.00125 | 0.00127 |
| Johri | 1.00E-06 | 1.00E+05 | 0.00394 | 0.00476 | 0.00237 | 0.00226 | 0.00224 |

**Table S9.** The ratio of diversity in neutral regions ($\pi_{neu}$) compared for *D. melanogaster* simulations with listed parameter combinations, compared to under strict neutrality. "With maps" refers to neutral simulations with realistic heterogeneity of mutation and recombination rates modelled, while "no maps" refers to neutral simulations where mutation and recombination rates were constant. The deleterious DFE that we used to simulate background selection (BGS) is indicated by the study that estimated the DFE in the first column. Parameters of recurrent positive selection employed in our simulations are provided by the proportion of new mutations that are beneficial ($f_{pos}$) and the mean population-scaled strength of selection ($2N_{anc}\bar{s_a}$).

| BGS | $f_{pos}$ | $2N_{anc}\bar{s_a}$ | Constant | Expansion | Contraction | Bottleneck | Repbneck |
|---|---|---|---|---|---|---|---|
| None (no maps) | N/A | N/A | 1.02 | 1.02 | 1.02 | 1.02 | 1.03 |
| None (with maps) | N/A | N/A | 1.00 | 1.00 | 1.00 | 1.00 | 1.00 |
| Johri | 0 | N/A | 0.40 | 0.48 | 0.30 | 0.48 | 0.54 |
| Johri | 0.0002 | 20 | 0.40 | 0.48 | 0.31 | 0.48 | 0.53 |
| Johri | 0.0002 | 250 | 0.35 | 0.37 | 0.27 | 0.46 | 0.51 |
| Johri | 0.005 | 20 | 0.39 | 0.46 | 0.30 | 0.48 | 0.53 |
| Johri | 0.005 | 250 | 0.12 | 0.09 | 0.12 | 0.25 | 0.28 |
| Johri | 1.00E-06 | 1.00E+05 | 0.34 | 0.33 | 0.27 | 0.45 | 0.50 |

**Table S10.** Alpha values (estimated proportion of beneficial substitutions at nonsynonymous sites) for *D. melanogaster* simulations with listed parameter combinations. Alpha at nonsynonymous sites was estimated from simulations by assuming that 25% of all coding sites are synonymous, using equation 2. The deleterious DFE that we used to simulate background selection (BGS) is indicated by the study that estimated the DFE in the first column. Parameters of recurrent positive selection employed in our simulations are provided by the proportion of new mutations that are beneficial ($f_{pos}$) and the mean population-scaled strength of selection ($2N_{anc}\bar{s_a}$).

| BGS | $f_{pos}$ | $2N_{anc}\bar{s_a}$ | Constant | Expansion | Contraction | Bottleneck | Repbneck |
|---|---|---|---|---|---|---|---|
| Johri | 0.0002 | 20 | 0.009 | 0.010 | 0.008 | 0.006 | 0.005 |
| Johri | 0.0002 | 250 | 0.093 | 0.111 | 0.084 | 0.066 | 0.060 |
| Johri | 0.005 | 20 | 0.178 | 0.189 | 0.166 | 0.128 | 0.114 |
| Johri | 0.005 | 250 | 0.447 | 0.466 | 0.438 | 0.416 | 0.403 |
| Johri | 1.00E-06 | 1.00E+05 | 0.011 | 0.015 | 0.010 | 0.008 | 0.007 |

**Table S11.** Divergence values in neutral intergenic/intronic regions for *D. melanogaster* simulations with listed parameter combinations. "With maps" refers to neutral simulations with realistic heterogeneity of mutation and recombination rates modelled, while "no maps" refers to neutral simulations where mutation and recombination rates were constant. The deleterious DFE that we used to simulate background selection (BGS) is indicated by the study that estimated the DFE in the first column. Parameters of recurrent positive selection employed in our simulations are provided by the proportion of new mutations that are beneficial ($f_{pos}$) and the mean population-scaled strength of selection ($2N_{anc}\bar{s_a}$).

| BGS | $f_{pos}$ | $2N_{anc}\bar{s_a}$ | Constant | Expansion | Contraction | Bottleneck | Repbneck |
|---|---|---|---|---|---|---|---|
| None (no maps) | N/A | N/A | 0.0119 | 0.0136 | 0.0168 | 0.0220 | 0.0288 |
| None (with maps) | N/A | N/A | 0.0117 | 0.0133 | 0.0164 | 0.0216 | 0.0282 |
| Johri | 0 | N/A | 0.0118 | 0.0135 | 0.0164 | 0.0175 | 0.0235 |
| Johri | 0.0002 | 20 | 0.0118 | 0.0135 | 0.0163 | 0.0174 | 0.0233 |
| Johri | 0.0002 | 250 | 0.0118 | 0.0139 | 0.0160 | 0.0169 | 0.0227 |
| Johri | 0.005 | 20 | 0.0118 | 0.0136 | 0.0162 | 0.0173 | 0.0233 |
| Johri | 0.005 | 250 | 0.0118 | 0.0148 | 0.0150 | 0.0149 | 0.0208 |
| Johri | 1.00E-06 | 1.00E+05 | 0.0118 | 0.0141 | 0.0160 | 0.0168 | 0.0227 |

**Table S12.** Divergence values in coding regions for *D. melanogaster* simulations with listed parameter combinations. "With maps" refers to neutral simulations with realistic heterogeneity of mutation and recombination rates modelled, while "no maps" refers to neutral simulations where mutation and recombination rates were constant. The deleterious DFE that we used to simulate background selection (BGS) is indicated by the study that estimated the DFE in the first column. Parameters of recurrent positive selection employed in our simulations are provided by the proportion of new mutations that are beneficial ($f_{pos}$) and the mean population-scaled strength of selection ($2N_{anc}\bar{s_a}$).

| BGS | $f_{pos}$ | $2N_{anc}\bar{s_a}$ | Constant | Expansion | Contraction | Bottleneck | Repbneck |
|---|---|---|---|---|---|---|---|
| None (no maps) | 0 | N/A | 0.0120 | 0.0136 | 0.0168 | 0.0221 | 0.0289 |
| None (with maps) | 0 | N/A | 0.0118 | 0.0134 | 0.0165 | 0.0217 | 0.0284 |
| Johri | 0 | N/A | 0.0050 | 0.0058 | 0.0071 | 0.0081 | 0.0113 |
| Johri | 0.0002 | 20 | 0.0051 | 0.0058 | 0.0071 | 0.0081 | 0.0113 |
| Johri | 0.0002 | 250 | 0.0057 | 0.0068 | 0.0078 | 0.0085 | 0.0118 |
| Johri | 0.005 | 20 | 0.0056 | 0.0065 | 0.0078 | 0.0087 | 0.0120 |
| Johri | 0.005 | 250 | 0.0113 | 0.0145 | 0.0141 | 0.0138 | 0.0190 |
| Johri | 1.00E-06 | 1.00E+05 | 0.0055 | 0.0065 | 0.0074 | 0.0082 | 0.0115 |

**Table S13.** Ratio of divergence values in coding/neutral regions for *D. melanogaster* simulations with listed parameter combinations. "With maps" refers to neutral simulations with realistic heterogeneity of mutation and recombination rates modelled, while "no maps" refers to neutral simulations where mutation and recombination rates were constant. The deleterious DFE that we used to simulate background selection (BGS) is indicated by the study that estimated the DFE in the first column. Parameters of recurrent positive selection employed in our simulations are provided by the proportion of new mutations that are beneficial ($f_{pos}$) and the mean population-scaled strength of selection ($2N_{anc}\bar{s_a}$).

| BGS | $f_{pos}$ | $2N_{anc}\bar{s_a}$ | Constant | Expansion | Contraction | Bottleneck | Repbneck |
|---|---|---|---|---|---|---|---|
| None (no maps) | 0 | N/A | 1.01 | 1.00 | 1.00 | 1.00 | 1.00 |
| None (with maps) | 0 | N/A | 1.01 | 1.00 | 1.01 | 1.01 | 1.01 |
| Johri | 0 | N/A | 0.43 | 0.43 | 0.43 | 0.46 | 0.48 |
| Johri | 0.0002 | 20 | 0.43 | 0.43 | 0.43 | 0.46 | 0.49 |
| Johri | 0.0002 | 250 | 0.48 | 0.49 | 0.48 | 0.50 | 0.52 |
| Johri | 0.005 | 20 | 0.48 | 0.48 | 0.48 | 0.50 | 0.52 |
| Johri | 0.005 | 250 | 0.96 | 0.98 | 0.94 | 0.92 | 0.91 |
| Johri | 1.00E-06 | 1.00E+05 | 0.46 | 0.46 | 0.47 | 0.49 | 0.51 |

**Table S14.** Nucleotide diversity in neutral intergenic/intronic regions ($\pi_{neu}$) and its value relative to the theoretical expectation of nucleotide diversity under neutrality ($4N\mu = 0.012$; see Methods and eqn. 3) for simulations of different scaling factors and length of simulated genome after $10N$ generations burn-in with constant population size. All simulations had 50% of new genome-wide mutations experiencing selection according to the DFE from Johri *et al.* (2020). Note, that the 15 MB, 100X simulations did not finish the burn-in within 86 hours.

| | | Length of simulated chromosome (L) | | | | | |
|---|---|---|---|---|---|---|---|
| | Scaling Factor | 500 kb | 1 Mb | 2 Mb | 3 Mb | 5 Mb | 15 Mb |
| $\pi_{neu}$ | 100X | 0.0102 | 0.0097 | 0.0091 | 0.0085 | 0.0078 | *DNF* |
| | 200X | 0.0096 | 0.0091 | 0.0083 | 0.0075 | 0.0062 | 0.0023 |
| | 300X | 0.0093 | 0.0086 | 0.0075 | 0.0066 | 0.0048 | 0.0011 |
| $\frac{\pi_{neu}}{0.012}$ | 100X | 0.85 | 0.81 | 0.76 | 0.71 | 0.65 | *DNF* |
| | 200X | 0.80 | 0.75 | 0.69 | 0.62 | 0.52 | 0.19 |
| | 300X | 0.78 | 0.72 | 0.62 | 0.55 | 0.40 | 0.09 |

**Table S15.** Mean counts of positively selected mutations that reached fixation in most recent 2N and 0.2N generations before sampling, both for all beneficial mutations and for strongly selected (s>0.05) beneficial mutations for human simulations. The deleterious DFE that we used to simulate background selection (BGS) is indicated by the study that estimated the DFE in the first column. Parameters of recurrent positive selection employed in our simulations are provided by the proportion of new mutations that are beneficial ($f_{pos}$) and the mean population-scaled strength of selection ($2N_{anc}\bar{s_a}$).

| | | | Fixations within last 2N generations | | Last 0.2N generations | |
|---|---|---|---|---|---|---|
| BGS | $f_{pos}$ | $2N_{anc}\bar{s_a}$ | All beneficial fixations | Beneficial fixations with s>0.05 | All beneficial fixations subs | Beneficial fixations of s>0.05 |
| Huber | 0.001 | 10 | 22.8 | 0 | 2.5 | 0 |
| Huber | 0.001 | 100 | 205.1 | 79.4 | 27 | 9.2 |
| Huber | 0.001 | 500 | 781.5 | 744.2 | 110 | 104.7 |
| Huber | 0.001 | 1000 | 1132.1 | 1118.8 | 158.8 | 157 |
| Huber | 0.01 | 10 | 209.1 | 0 | 30.2 | 0 |
| Huber | 0.01 | 100 | 1574 | 728.4 | 229.8 | 106.2 |
| Huber | 0.01 | 500 | 3208.3 | 3125.5 | 458.6 | 446.6 |
| Huber | 0.01 | 1000 | 3608.6 | 3589.1 | 518.2 | 515.3 |

**Table S16.** Mean counts of positively selected mutations that reached fixation in most recent 2N and 0.2N generations before sampling, both for all beneficial mutations and for strongly selected (s>0.05) beneficial mutations for *D. melanogaster* simulations. The deleterious DFE that we used to simulate background selection (BGS) is indicated by the study that estimated the DFE in the first column. Parameters of recurrent positive selection employed in our simulations are provided by the proportion of new mutations that are beneficial ($f_{pos}$) and the mean population-scaled strength of selection ($2N_{anc}\bar{s_a}$).

| | | | Fixations within last 2N generations | | Last 0.2N generations | |
|---|---|---|---|---|---|---|
| BGS | $f_{pos}$ | $2N_{anc}\bar{s_a}$ | All beneficial fixations | Beneficial fixations with s>0.05 | All beneficial fixations subs | Beneficial fixations of s>0.05 |
| Johri | 0.0002 | 20 | 30.1 | 0 | 3.4 | 0 |
| Johri | 0.0002 | 250 | 411.3 | 257.4 | 39.4 | 24.6 |
| Johri | 0.005 | 20 | 777.9 | 0 | 77.5 | 0 |
| Johri | 0.005 | 250 | 6154.4 | 4260.7 | 621.6 | 436.2 |
| Johri | 1.00E-06 | 1.00E+05 | 48.8 | 48.8 | 5.5 | 5.5 |
